# An electron transfer competent structural ensemble of membrane-bound cytochrome P450 1A1 and cytochrome P450 oxidoreductase

**DOI:** 10.1101/2020.06.12.149112

**Authors:** Goutam Mukherjee, Prajwal P. Nandekar, Rebecca C. Wade

## Abstract

Cytochrome P450 (CYP) heme monooxygenases require two electrons for their catalytic cycle. For mammalian microsomal CYPs, key enzymes for xenobiotic metabolism and steroidogenesis and important drug targets and biocatalysts, the electrons are transferred by NADPH-cytochrome P450 oxidoreductase (CPR). No structure of a mammalian CYP-CPR complex has been solved experimentally, hindering understanding of the determinants of electron transfer (ET), which is often rate-limiting for CYP reactions. Here, we investigated the interactions between membrane-bound CYP 1A1, an antitumor drug target, and CPR by a multiresolution computational approach. We find that upon binding to CPR, the CYP 1A1 catalytic domain becomes less embedded in the membrane and reorients, indicating that CPR may affect ligand passage to the CYP active site. Despite the constraints imposed by membrane binding, we identify several arrangements of CPR around CYP 1A1 that are compatible with ET. In the complexes, the interactions of the CPR FMN domain with the proximal side of CYP 1A1 are supplemented by more transient interactions of the CPR NADP domain with the distal side of CYP 1A1. Computed ET rates and pathways agree well with available experimental data and suggest why the CYP-CPR ET rates are low compared to those of soluble bacterial CYPs.

## Introduction

CYP enzymes constitute the cardinal xenobiotic-metabolizing enzyme superfamily and they are of considerable interest for the synthesis of novel drugs and drug metabolites, targeted cancer gene therapy, biosensor design and bioremediation.^1–3^ CYPs are widely used as biocatalysts for regio- and enantioselective C-H hydroxylation reactions. CYPs are heme monooxygenases that activate one molecule of dioxygen and oxidize the substrate by inserting one oxygen atom to form the product and reducing the other oxygen atom to water. Thus, drugs are transformed into more polar metabolites during the process of drug metabolism.^4^

In the catalytic cycle, CYP requires two electrons to be transferred sequentially from a redox partner to its heme cofactor (HEME), and the ET steps have been observed to be rate-limiting for the reaction in many cases.^5^ For the microsomal CYPs, the redox partner is CPR, a flavoprotein containing three cofactor binding domains: the FMN domain with the flavin mononucleotide (FMN) cofactor, the FAD domain with the flavin adenine dinucleotide (FAD) cofactor, and the NADP domain with the nicotinamide adenine dinucleotide phosphate (NADP) cofactor. Besides these three domains, CPR also contains an amino-terminal membrane binding domain, which serves as a nonspecific membrane anchor, and a flexible tether region that connects this domain and the FMN domains^6^. The membrane binding domain is necessary for efficient ET from CPR to CYP^6^ and its interactions with the membrane have been found to depend on cofactor binding and the oxidation state of CPR.^7^ Cytochrome b5 (cyt b5) is another natural redox partner for some CYPs and can also allosterically modulate CYP catalysis.^8–11^

All mammalian CYPs have three domains: a globular catalytic HEME-containing domain, an N-terminal transmembrane α-helical domain, and a flexible linker region connecting these two domains. In humans, both the CYP and the CPR are anchored in the endoplasmic reticulum (ER) membrane by a transmembrane helix (TM-helix). The N-terminal region of the CYPs not only anchors the protein in the ER membrane, but also influences their localization to different microdomains of the membrane.^12^

Information about the orientation of the globular domain of CYPs, alone or in CYP-CPR/cyt b5 complexes in the membrane, has been gleaned from experiments such as linear dichroism measurements in Nanodiscs,^13,14^ atomic force microscopy and antibody mapping.^15^ To date, there is no experimentally determined structure of a full CYP-CPR/cyt b5 complex available, although structures of the individual globular domains of a number of mammalian CYPs and CPRs have been determined by crystallography after removal of the N-terminal transmembrane domain to facilitate expression and crystallization.^2^ Various modeling and simulation procedures have been developed to predict the orientation of full-length CYPs in phospholipid bilayers.^16–22^ Notably, *T. brucei* CYP 51 simulated^20^ prior to the determination in 2014 of the crystal structure of yeast CYP 51,^23^ the only eukaryotic CYP for which a full-length structure has been determined experimentally, indicated a very similar orientation of the protein with respect to the membrane. Experiments, such as solid-state NMR and mutagenesis, indicate that the association between a CYP and its CPR/cyt b5 redox partner is driven by electrostatic and hydrophobic interactions with positively charged residues on the proximal side of CYP interacting with negatively charged residues on the CPR FMN domain or cyt b5^24–29^.

We initially took the crystal structure of *Bacillus megaterium* cytochrome P450 BM3 (P450_BM3_)^30^, a non-stoichiometric complex of two CYP HEME domains and an FMN domain, as a template, for modeling the CYP 1A1-CPR complex but, despite extensive MD simulation, we did not succeed in obtaining ET-competent complexes. Indeed, considering the P450_BM3_ structure as a template for modelling mammalian CYP-CPR interactions has four major drawbacks: (i) The crystal structure of P450_BM3_ (PDB ID: 1BVY) is not consistent with measured ET rates due to the high redox center separation distance; (ii) The low sequence identity between P450_BM3_ and CYP/CPR of about 30% for the CYP and FMN domains; (iii) Unlike mammalian CYPs, the dimeric form of P450_BM3_ is catalytically functional and ET from FMN to HEME is proposed to occur in a trans fashion;^31,32^ (iv) Mammalian CYP and CPR are membrane-anchored proteins, whereas P450_BM3_ is not and the membrane attachment may affect the binding of CPR to the CYP. Hence, it is preferable to employ a *de novo* protocol rather than a template-based method for modelling full-length CYP-CPR interactions in a membrane.

Here, we have carried out multiresolution simulations with the aim of achieving a detailed understanding of the structure and dynamics of the membrane-associated full-length CYP 1A1-CPR complex, and consequently of the determinants of the ET process between CYP 1A1 and CPR. CYP 1A1 is an extrahepatic human drug target protein that plays an important role in the bioactivation of procarcinogens and detoxification of carcinogens.^33,34^ Many of its substrates, such as benzo[o]pyrene, a carcinogen found in tobacco smoke, are polycyclic aromatic hydrocarbons. The standard reference substrate for CYP 1A1 is 7-ethoxyresorufin, which is O-dealkylated by the enzyme. Constitutive expression of CYP 1A1 is low but can be induced in response to environmental contaminants, for example in smokers’ epithelial lung cells. Moreover, engineered CYP 1A1 has potential as a biocatalyst.^35,36^ Notably, CYP 1A1-CPR fusion proteins have been made to improve ET rates. A fusion between rat CYP 1A1 and rat CPR was found to exhibit four times higher monooxygenase activity than rat CYP 1A1 alone,^35^ whereas a fusion between rat CYP 1A1 and yeast CPR had a similar ET rate to P450_BM3_.^36^ Hence, a detailed atomic level understanding of CYP 1A1-CPR interactions and ET kinetics would provide the basis for the design of drugs targeting CYP 1A1, as well as the exploitation of CYP 1A1 as a biocatalyst. We therefore here describe the building and simulation of the full-length CYP 1A1–CPR complex in a 1-palmitoyl-2-oleoyl-sn-glycero-3-phosphocholine (POPC) bilayer. POPC was chosen because phosphatidylcholine is the main component in the mammalian ER membrane^37^ and because it provides a simple representation, which is often used in *in vitro* studies of CYPs, of the disordered regions of the ER where CYP 1A1 has been observed to localize.^12,38^

## Results and Discussion

### Electrostatic steering of the CYP 1A1 and the CPR FMN domains leads to encounter complex formation

The multiresolution computational method used to model and simulate the membrane-associated full-length CYP 1A1-CPR complex is depicted in **Fig. 1**, (further details are given in **Methods, Supplementary Methods** and **Fig. S1)**. The Brownian dynamics (BD) docking method allows generation of diffusional encounter complexes that satisfy biochemical constraints. It has been successfully applied to predict protein-protein complexes,^39^ including those between cytochrome P450s and their electron transfer partners. ^39–41^ For example, the orientation of putidaredoxin in the predicted complex with cytochrome P450cam^42^ was similar (about 15° rotated) to that in the crystal structure of the complex published afterwards by Tripathi et al.^43^ The six representative encounter complexes of the globular domains of CYP 1A1 and CPR generated by BD rigid-body docking simulations and selected for subsequent molecular dynamics (MD) simulations are listed in **Table 1 (**all encounter complexes are listed in the **Supplementary MS Excel sheet** and **Table S1)**. The representative encounter complexes and their expected position with respect to the membrane are shown in **Fig. S2** and one of these complexes is shown in **Fig 2a**. The interfacial residues in these complexes are listed in **Table S2**.

**Table 1:** Evolution of the CYP 1A1-FMN domain interface and electron transfer (ET) rates of the complexes along the steps of the multiresolution simulation procedure. Observed redox center distances (D_Fe-N5_), center-to-center distances between the CYP and FMN domains of CPR (D_CYP-FMN domain_), θ angle, and calculated ET rates are given for six complexes.

| Sys-tem | Energy minimized complexes generated by BD simulation |  |  |  | Complexes from MD simulation in explicit aqueous solution without a membrane<sup>s</sup> |  |  |  | Complexes from MD simulations in membrane-bound form<sup>&</sup> |  |  |  |
| --- | --- | --- | --- | --- | --- | --- | --- | --- | --- | --- | --- | --- |
| | $D_{\text{Fe-N5}}$ (Å) | $D_{\text{CYP-FMN domain}}$ (Å) | $\theta$ (°) | $k_{\text{ET}}$ (s<sup>-1</sup>) | $D_{\text{Fe-N5}}$ (Å) | $D_{\text{CYP-FMN domain}}$ (Å) | $\theta$ (°) | $k_{\text{ET}}$ (s<sup>-1</sup>)<sup>*</sup> | $D_{\text{Fe-N5}}$ (Å) | $D_{\text{CYP-FMN domain}}$ (Å) | $\theta$ (°) | $k_{\text{ET}}$ (s<sup>-1</sup>) |
| A4 | 19.2 | 34.6 | 53.5 | 0.002 | 19.4±0.4 | 35.7±0.3 | 107.6±3.2 | 0.002 | -- | -- | -- | -- |
| B7 | 16.3 | 37.3 | 89.6 | 0.001 | 21.7±0.9 | 42.7±0.7 | 124.2±6.4 | 0.000 | -- | -- | -- | -- |
| C2 | 18.4 | 37.4 | 94.9 | 0.001 | 17.2±0.3 | 36.1±0.3 | 92.8±2.6 | 0.004 | 15.6±0.3 | 34.6±0.2 | 95.7±4.0 | -- |
| C3 | 17.5 | 35.7 | 99.0 | 0.005 | 15.5±0.3 | 33.8±0.2 | 98.3±2.0 | 0.013 | 15.5±0.4 | 33.7±0.2 | 99.1±1.7 | 0.2±0.5 |
| D2 | 15.4 | 37.9 | 70.4 | 0.000 | 14.5±0.3 | 36.5±0.3 | 63.1±2.5 | 1.882 | 14.7±0.4 | 35.6±0.4 | 62.0±3.1 | 8.1±16.0 |
| D2'<sup>#</sup> |  |  |  |  |  |  |  |  | *16.3±0.3* | *33.0±0.2* | *75.1±2.6* | *2.4±2.6* |
| D3 | 20.2 | 35.2 | 95.3 | 0.000 | 20.0±0.6 | 36.4±0.4 | 104.3±2.9 | 0.000 | -- | -- | -- | -- |

<sup>\$</sup>Average over last 20 ns with 2 ps intervals; <sup>&</sup>Average over last 50 ns with 100 ps intervals; <sup>\*</sup>Last frame. <sup>#</sup>Values in italics are for the second set of simulations of the D2 complex (D2').
*(a) Weakening of CYP 1A1-membrane interactions and reorientation of CYP globular domain*

**Figure 1:**
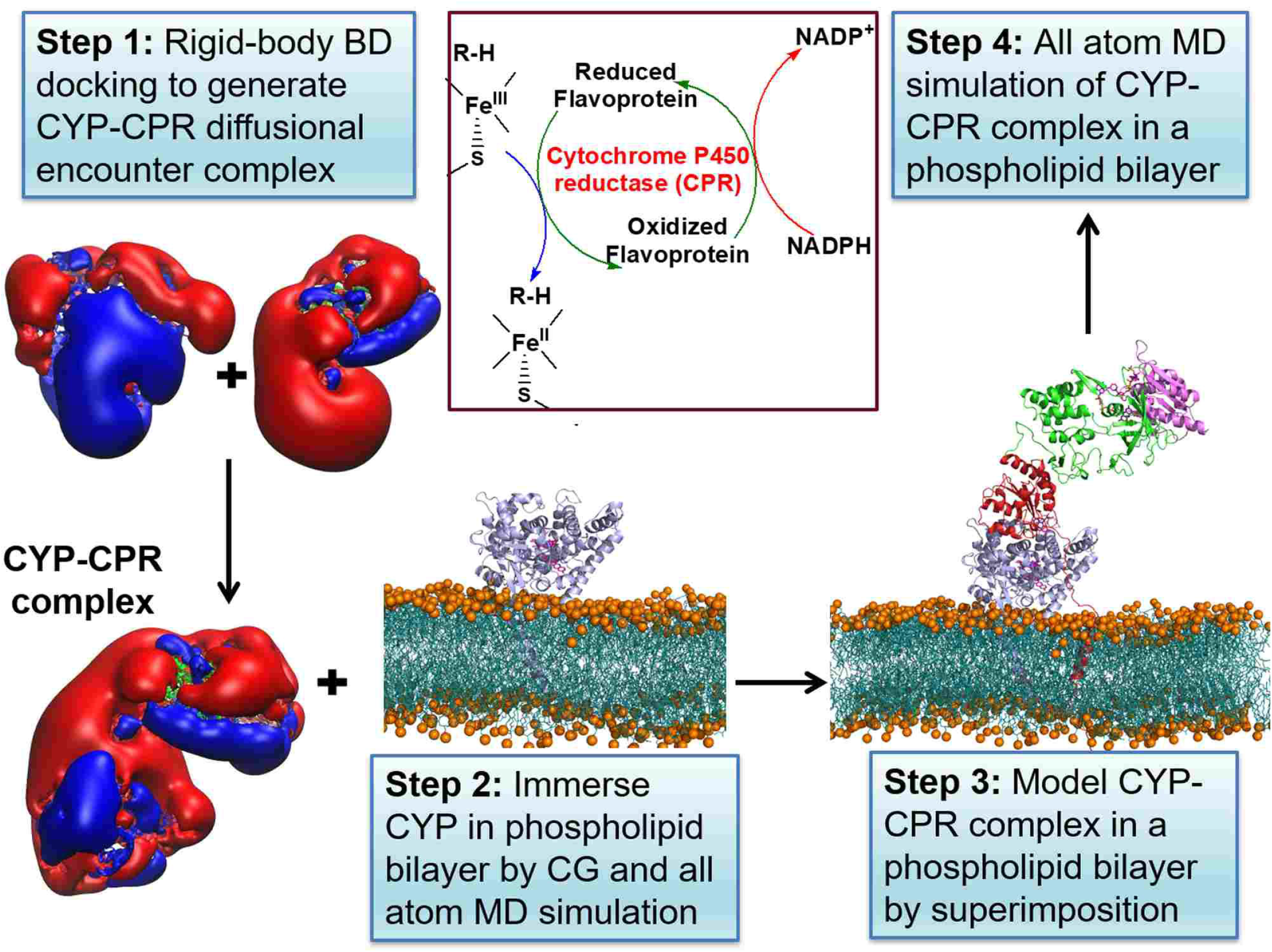
Diagram of the procedure to build and simulate a model of a mammalian CYP-CPR complex in a membrane. The formation of the CYP-CPR complex is necessary for the transfer of electrons to the CYP active site in the CYP catalytic cycle, as indicated in the schematic cycle. Step 1: Brownian dynamics (BD) rigid-body docking of CYP and CPR globular domains. Molecular electrostatic isopotential contours at ±1 kT/e show a highly positive (blue) patch on the proximal face of CYP and a highly negative (red) patch on the CPR that interact complementarily in the docked complexes. Step 2: Coarse-grained (CG) and all-atom molecular dynamics (MD) simulation of CYP (blue cartoon representation) in a phospholipid bilayer (cyan with orange spheres representing phosphorous atoms). Step 3: Relaxation of the BD docked complexes by MD simulation in aqueous solution followed by superimposition on the CYP in the bilayer (with CPR shown in cartoon representation colored by domain (FMN: red, FAD: green, NAD: pink)). Step 4: Atomic detail MD simulation of the CYP-CPR complexes in a phospholipid bilayer.

**Figure 2:**
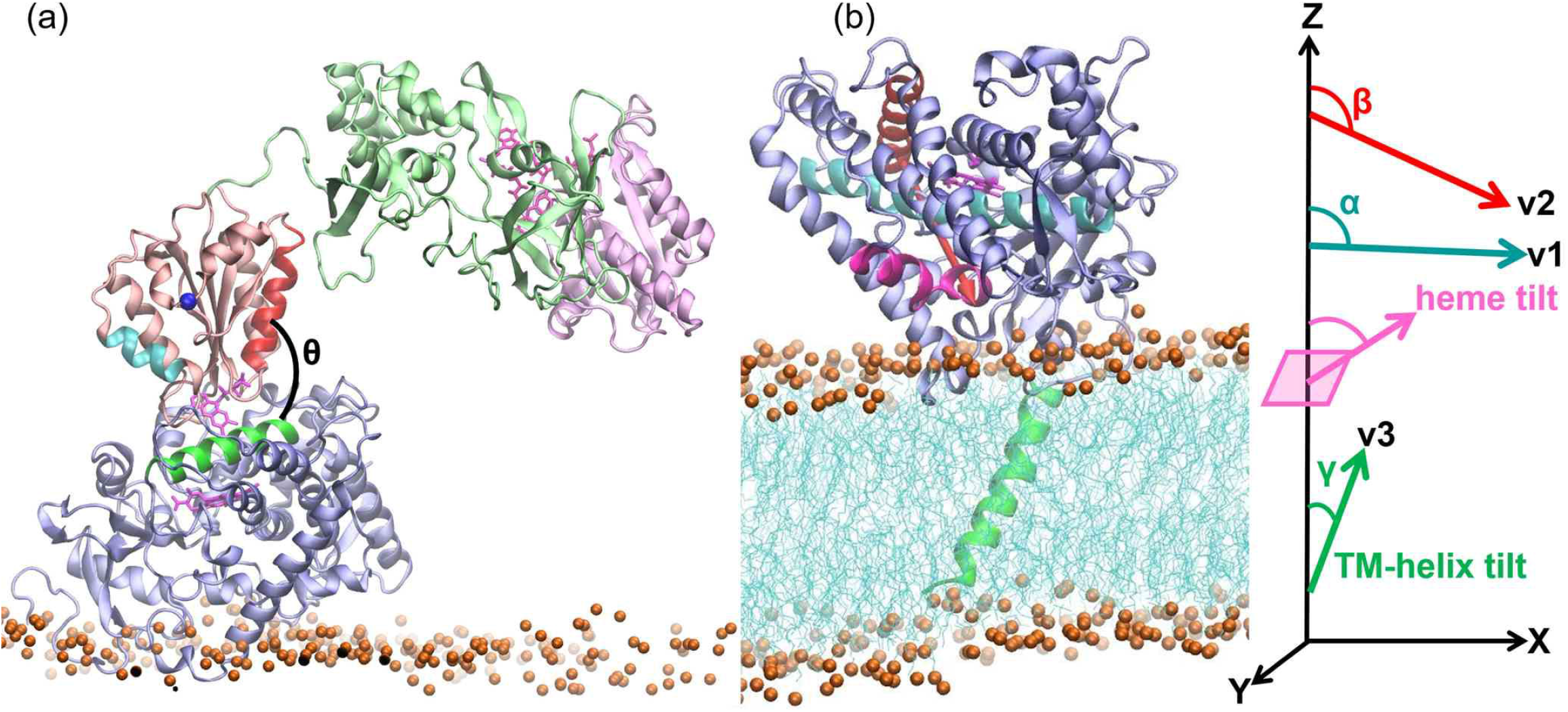
A CYP 1A1-CPR encounter complex obtained from the rigid-body BD docking simulations is shown superimposed on the structure of CYP 1A1 in a membrane bilayer obtained from MD simulations which is shown together with the definitions of the angles defining the arrangement of the proteins. A CYP 1A1-CPR encounter complex (D2) obtained from the rigid-body BD docking simulations is shown in (a) superimposed on the structure of CYP 1A1 in a membrane bilayer obtained from MD simulations shown in (b). The proteins are shown in cartoon representation (CYP globular domain: lilac, TM-helix: green; CPR FMN domain: salmon, FAD domain: pale green, NADP domain: pale pink), cofactors are shown in pink stick representation, and the bilayer is shown in cyan lines with orange spheres representing the phosphorous atoms. The angles defining the arrangement of the proteins are shown: θ is the angle between the CYP C-helix (green) and the FMN domain α1-helix (residues 91-105; red); the FMN domain α3-helix (150-158; cyan) and N-terminal residue of FMN domain (blue sphere) are indicated; the heme tilt angle is the angle between the heme plane and the z-axis perpendicular to the membrane plane; α, β and γ are the angles between the z-axis and the vectors v1, along the I-helix (cyan), v2, orthogonal to v1 and connecting the C-helix (red) and the F-helix (magenta); and v3 along the TM-helix (residues 7-26; green).

In the encounter complexes, the globular domains of CYP 1A1 and CPR dock specifically due to complementary electrostatic interactions, and the proximal face of CYP 1A1 and the outer FMN-binding loop interact (**Fig. 1**). The FMN domain binding mode differs from that in the crystal structure of the P450_BM3_ CYP-FMN domain complex.^30^ In the encounter complexes, both the α1- and α3-helices of the FMN domain contribute to the interface whereas, in P450_BM3_, it is mainly the α1-helix of the FMN domain. The distance between the redox centers, D_Fe-N5_, in the six representative encounter complexes ranges from 15.4-20.2 Å and is shorter than the distance (23.6 Å) observed in the crystal structure of P450_BM3_ even though the distance between the centers of mass of the globular domain of CYP 1A1 and the FMN domain, D_CYP-FMN domain_, is higher in the representative diffusional encounter complexes (34.6-37.9 Å) than in the P450_BM3_ crystal structure (31.7 Å). (**Table 1**). The relative orientation of the FMN domain and the CYP globular domain can be characterized^44^ by the angle θ between the α_1_-helix of the FMN domain and the CYP C-helix (see **Fig. 2a**). Angle θ ranges from 54° to 99° in these 6 complexes, whereas in the P450_BM3_ crystal structure, it is 95°. Nevertheless, the binding face of CYP 1A1 in all these complexes is similar to that identified in mutagenesis studies of CYP 1A2-CPR^45^ and CYP 2B4-CPR binding.^27^ Next, we relaxed the BD rigid-body docked complexes by performing MD simulations.

### Structural relaxation of encounter complexes in “soluble” MD simulations resulted in three putative ET-competent complexes

All “soluble” MD simulations were performed for the globular domain of apo-CYP 1A1 and the FMN domain of CPR in a periodic box of aqueous solvent with initial structures taken from the six selected diffusional encounter complexes. The structures of the individual domains were well maintained throughout all the production simulations. The C_α_ atom root mean squared deviation (C_α_-RMSD) values of the CYP globular domain and the FMN domain were 2.0-2.6 Å and 1.2-1.4 Å, respectively (**Table S3**). The arrangement of the protein domains in the complexes relaxed during the simulations (**Fig. 3**). Three of the six encounter complexes were discarded after structural relaxation, either because they were no longer ET-competent (B7 and D3, see **Table 1**) or because of rearrangements in the hydrogen-bonding of the FMN cofactor (A4, see **Fig. S3**).

**Figure 3:**
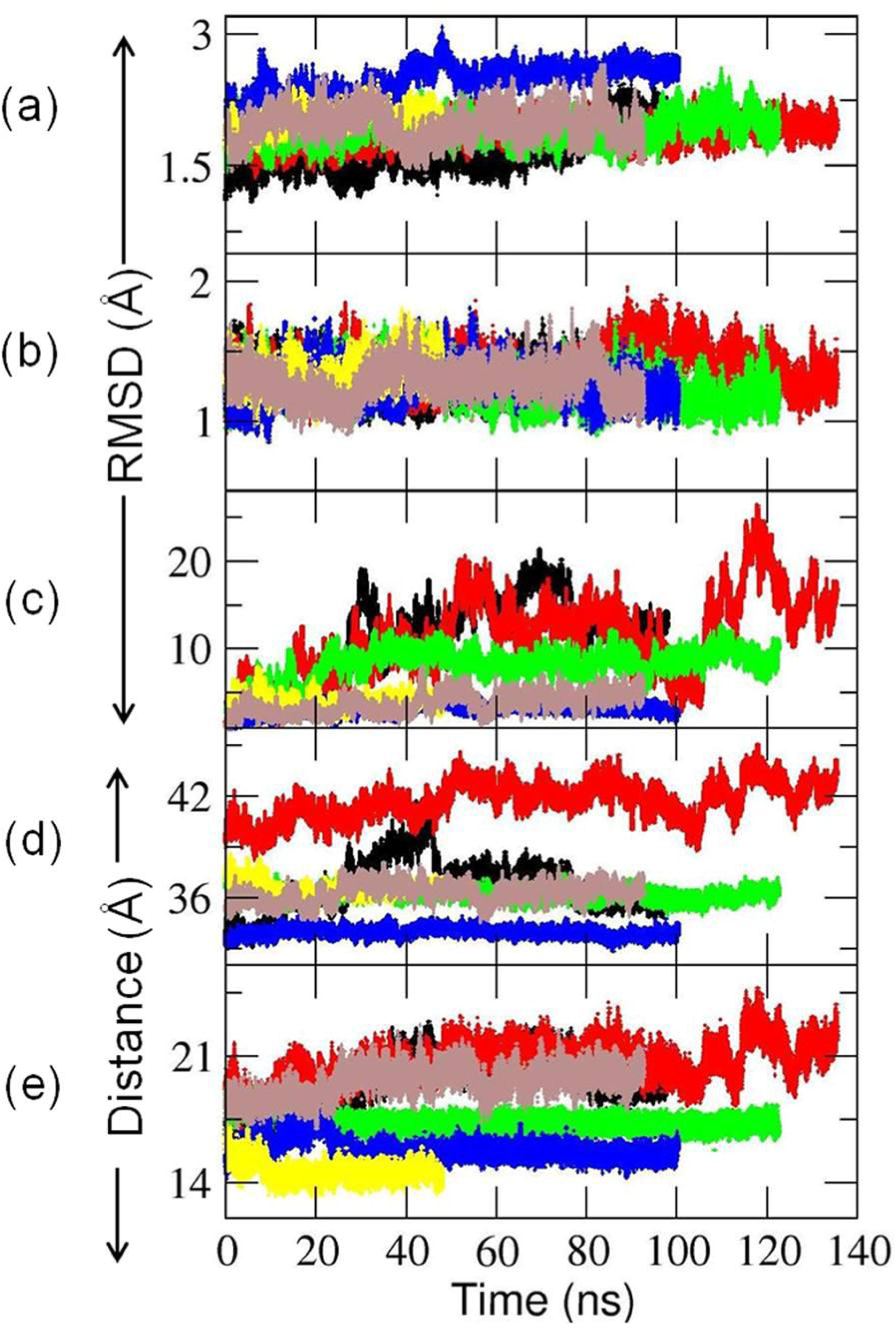
Evolution of the six CYP-FMN domain encounter complexes generated by BD rigid body docking during the MD simulations in aqueous solution (referred to as ‘soluble’ simulations). C_α_ atom root mean squared deviation (C_α_-RMSD) of (a) the globular domain of CYP 1A1, (b) the FMN domain of CPR, and (c) the FMN domain when the CYP domains of the complex were superimposed with respect to the initial frame. (d) Center-to-center distance between the CYP and the FMN domain, D_CYP-FMN domain_, and (e) the distance between the HEME:Fe and FMN:N5 atoms, D_Fe-N5_. RMSDs were calculated with respect to the initial energy minimized encounter complexes generated by BD simulation. Color scheme: A4: Black; B7: Red; C2: Green; C3: Blue; D2: yellow; D3: Brown.

For the remaining three encounter complexes (C2, C3 and D2), decreases in D_CYP-FMN domain_ and D_Fe-N5_ were observed during the simulations (**Table 1**), indicating the formation of tighter complexes with non-zero ET rates. The interface residues of CYP 1A1 remained similar to those in the initial BD docked complexes (**Table S2**). Next, these three refined complexes were used to build and simulate complete CYP-CPR complexes in the presence of a phospholipid bilayer.

### CYP-membrane interactions weaken in the presence of CPR and the CPR structure adopts a more compact form

All three models of membrane-bound CYP-CPR complexes were simulated for about 500 ns. In addition, a second simulation (D2’), with identical initial coordinates but with different initial assigned velocities, was run for the D2 system. The structures of the individual protein domains were well maintained as shown in C_α_-RMSD plots (**Fig. S4)** but there were rearrangements of the domains within CPR and of the proteins with respect to the membrane (**Table S4**).

#### a) Weakening of CYP 1A1-membrane interactions and reorientation of CYP globular domain

During the simulations, the axial distance between the centers of mass of the globular domain of CYP 1A1 and the membrane, D_CYP-mem_, increased by 3-8 Å (**Table S4**) indicating that, in the presence of CPR, CYP becomes less deeply embedded in the membrane. The protein-lipid interactions of CYP 1A1 are mainly hydrophobic in nature and formed by a completely immersed TM-helix and a partially immersed FG-loop (residues 229–245). During all the simulations except for C3, the axial distance between the centers of mass of the FG-loop and the membrane, D_FG-mem_, increased by 3-5 Å (**Table S4**), indicating a weakening of the peripheral CYP interactions with the membrane.

In the presence of CPR, the CYP 1A1 globular domain underwent an orientational rearrangement with respect to the CPR-free membrane-bound state as indicated by the changes in α, β and heme-tilt angles^16^ (**Fig. 2b**) along the simulations (**Fig. 4, Table S4**). Despite the almost identical starting orientations of CYP 1A1 in the membrane, the simulations converge to quite different orientations of the CYP globular domain due to the different binding modes of CPR to CYP 1A1. In the presence of CPR, the heme tilt angle dropped from ∼71° to around 40° for the C2 and D2 systems (**Table S4**). Thus, the interactions with CPR, resulting in an increase of the area of the FMN domain interface of up to ∼700 Å^2^, weakened the CYP-membrane interactions, and facilitated reorganization of the CYP with respect to the membrane.

**Figure 4:**
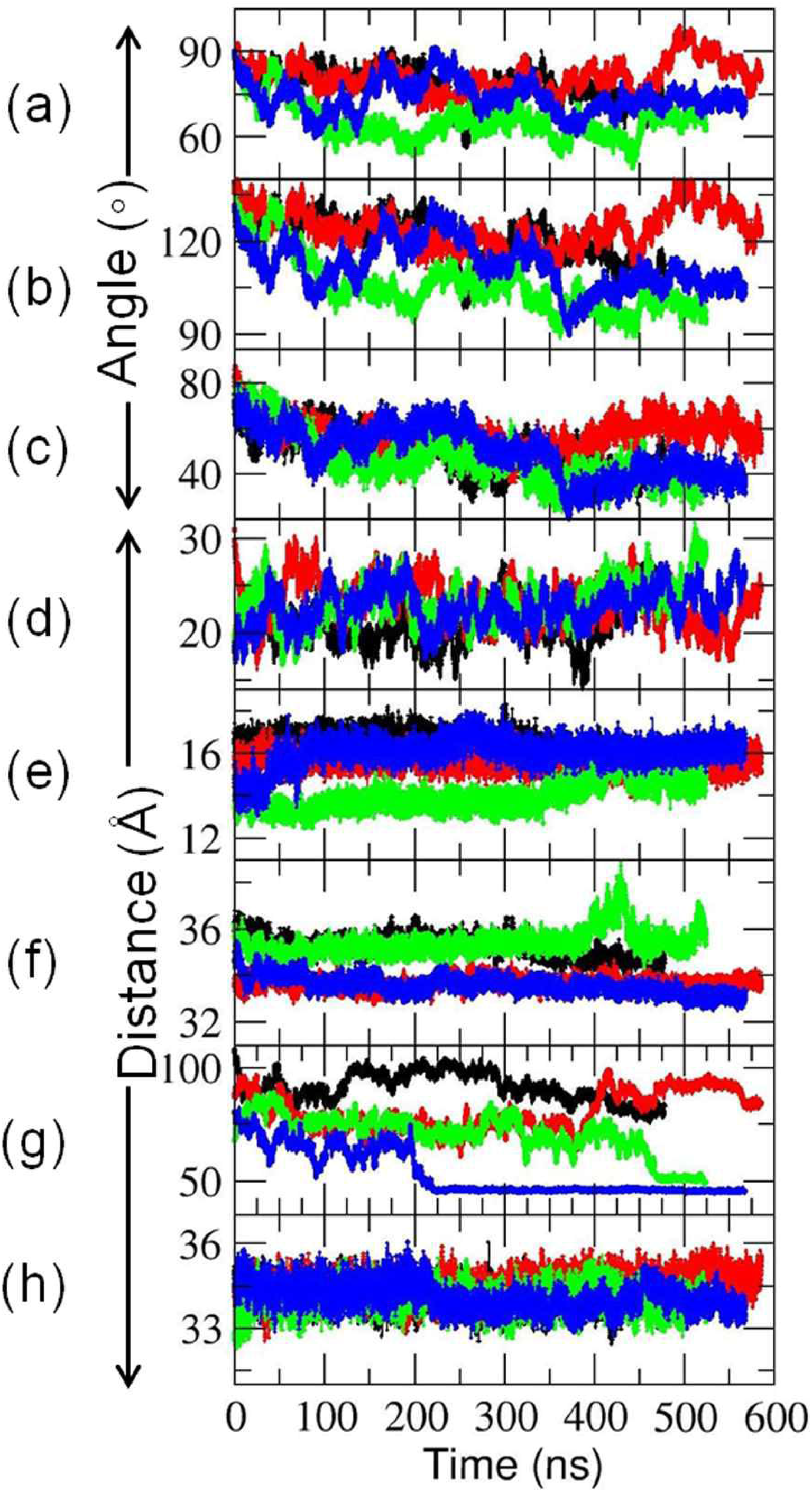
Observed changes in the position and configuration of the CYP-CPR complexes in the phospholipid bilayer during MD simulations. (a-c) Angles defining the CYP orientation in the membrane: (a) α, (b) β, and (c) heme tilt; (d): center of mass (CoM) distance of the FG loop to the CoM of the membrane, D_FG-mem_; (e) the redox center separation distance, D_Fe-N5_; (f-h): CoM distances between (f) CYP and FMN domains, D_CYP-FMN domain_ (g) CYP and NADP domains, D_CYP-NADP domain_ and (h) FAD and NADP domains, D_FAD-NADP domain_. The lengths of the simulations for C2 (black), C3 (red), D2 (green) and D2’ (blue) were 478, 585, 524 and 568 ns, respectively.

#### b) Rearrangement of the FMN domain of CPR with respect to the membrane

For CPR, a greater extent of rearrangement of the FMN domain with respect to the membrane was observed in the D2 and D2’ simulations as indicated by the large drop in D_FMN domain-mem_ (**Table S4**). This drop was accompanied by steep decreases of more than 43 Å in the axial distance, D_linker-mem,_ between the first atom of the FMN domain (the amide nitrogen of E66) and the center of mass (CoM) of the membrane for the D2 simulations. Thus, rearrangements of the highly flexible linker region between the TM-helix of CPR and the FMN domain affected the CPR orientation with respect to the membrane. However, this large rearrangement of the FMN domain towards the membrane had little effect on the interdomain distance, D_CYP-FMN domain_.

#### c) Rearrangement of the FAD and NADP domains with respect to the membrane results in a more compact structure of CPR

Due to the highly flexible linker hinge between the FMN and FAD domains of CPR, rearrangements of the FAD and NADP domains with respect to the membrane were observed (**Fig. 5 a-d, Table S4)**. This domain motion, as well as the initial binding mode of the encounter complexes, brought the FAD and NADP domains nearer to the globular domain of CYP 1A1 in the C2 and D2 systems (**Fig. 4**). For the D2’ simulation, the decrease in the FAD-NADP interdomain distance resulted in a more compact CPR structure, similar to that of the semi-open conformation in the crystal structure (PDB ID 3ES9 chain A) (**Fig. S5)**. In this structure, the center-to-center distance between the FMN and FAD cofactors was reduced to 44.7 Å, a distance intermediate between that of the closed form (PDB ID 3QE2) of 14.2 Å and that of the fully open form (PDB ID: 3ES9 chain B) of 61.3 Å, and too long for ET between them. This compaction of CPR enabled additional contacts to be formed between the NADP domain and the distal side of CYP 1A1.

**Figure 5:**
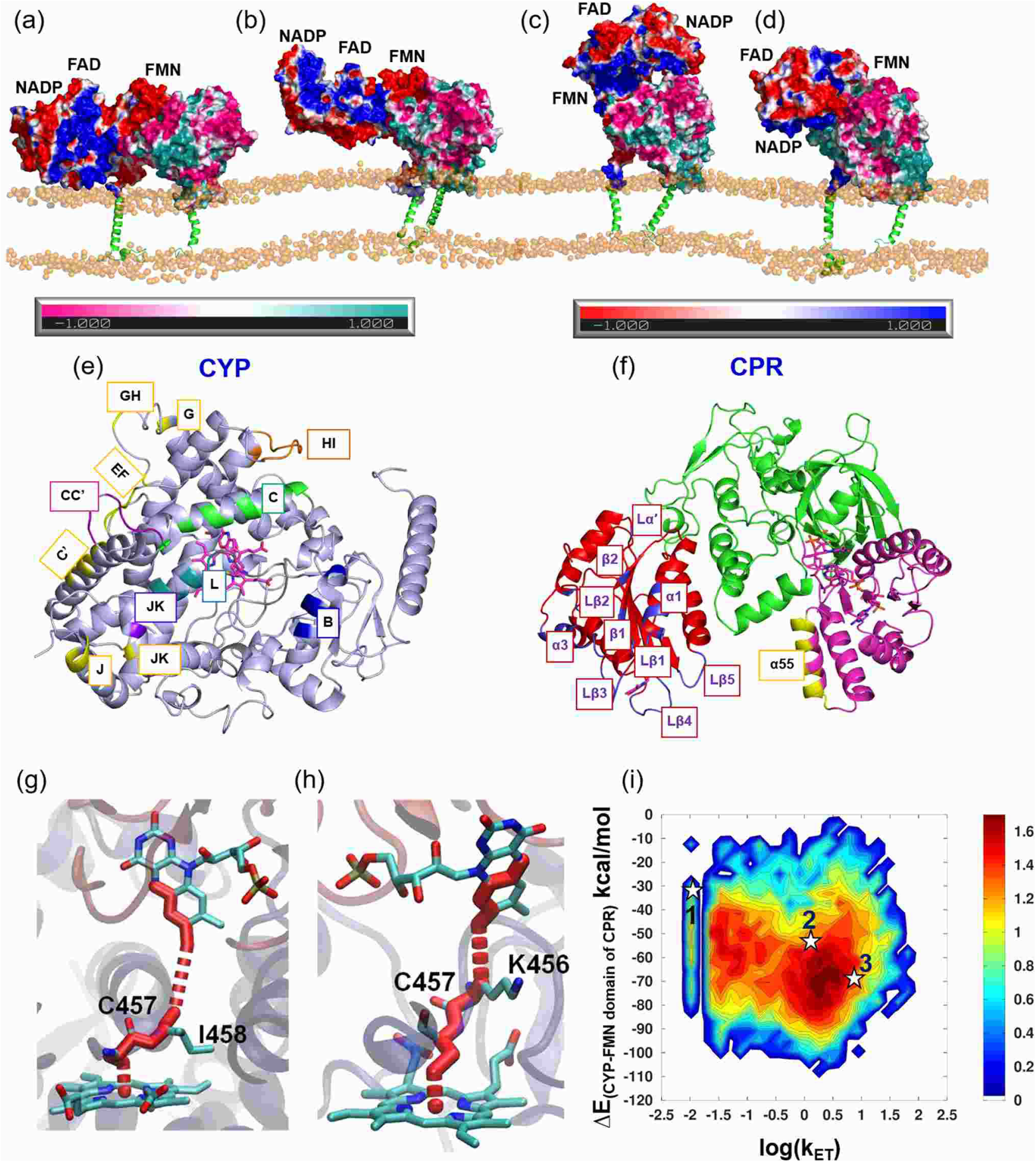
Structures, interactions and predicted ET pathways of four CYP-CPR complexes in a POPC bilayer obtained from MD simulations. (a) C-2, (b) C3, (c) D2 and (d) D2’. The proteins are shown in surface representation colored by electrostatic potential (positive: CYP (cyan) and CPR (blue); negative: CYP (pink) and CPR (red)) with the transmembrane helices in green cartoon representation and the phosphorous atoms of the lipids represented as orange spheres. (e,f) The interfaces in the model CYP 1A1-CPR complexes are highlighted by colors and labels on cartoon representations of (e) CYP 1A1 (light blue) and (f) CPR (with the FMN domain (red), the FAD domain (green) and the NADP domain (pink)); CYP interface colors: B-Helix: Blue; C-Helix: Green; CC’-Loop: Purple; JK-Loop: Violet; HI-Loop and I-Helix: Orange; Loop near HEME: Gray; L-Helix: Deep teal; NADP binding region: Yellow. The FMN domain interface is colored blue and the NADP domain interface is colored yellow. The interfacial residues are listed in Table S2. (g-i) Predicted ET pathways from the N5 of the FMN cofactor of CPR to Fe of the CYP 1A1 heme cofactor in complexes from (g) C3, D2, and (h) D2’ simulations; (i) 2D-histogram plot of the computed free energy of formation of the CYP-FMN domain complex vs predicted log(k_ET_) rates. Stars indicate complexes from the last frames of the simulations (1) C3, (2) D2 and (3) D2’ for which atomic coordinates are provided in ModelArchive.

### CYP 1A1 and CPR form stable complexes in the membrane

In all the simulations, the interdomain distance DCYP-FMN domain was stable at 33-36 Å. It remained longer than the interdomain distance in the crystal structure of P450_BM3_ (31.7 Å), possibly because the membrane prevents the two domains from coming as close as in the membrane-free P450_BM3_. Nevertheless, in all the CYP 1A1-CPR complexes, D_Fe-N5_ was within the range (14-16 Å) for biological ET (**Table S4**) and converged stably in all the simulations (**Fig. 4**).

No contacts between the TM-helices of CYP 1A1 and CPR were observed, which is consistent with the observations of Sundermann and Oostenbrink^46^ in their 10 ns simulation of a membrane-bound CYP 2D6-CPR complex. However, the computed interface contact area of the globular domain of CYP 1A1 and the CPR FMN domain in the model complexes was ∼2000 Å^2^ (**Table S4**), much higher than the interface area of about 240 Å^2^ observed in the simulation of a membrane-bound CYP 2D6-CPR complex,^46^ and greater that the interface area of many transient protein-protein complexes, which is usually is less than 1500 Å^2^.^47,48^ This large interface area is consistent with the measured binding affinity of CPR to rat CYP 1A2 (which has 68% identity to human CYP 1A1) of 47 nM.^45^ The CPR NADP domain interactions with the distal side of CYP create an additional contact interface. The high contact area of the CYP 1A1-CPR complexes indicates the formation of a rather strongly bound transient complex between the proteins.

### The interface between CYP 1A1 and the CPR FMN domain mostly has similar residues to other CYP-CPR complexes but shows differences to the P450_BM3_ CYP-FMN domain interface

The CYP-FMN domain interactions involve charge pairing of the positively charged proximal side of CYP 1A1 with the negatively charged surface of the FMN domain surrounding the cofactor. These electrostatic interactions are complemented by van der Waals and hydrophobic interactions. The interface residues in CYP 1A1 are located on the B, C and L-helices and the JK-loop and the loop structure near the HEME (**Table S2** and **Fig. 5e**). The interface residues of the CPR FMN domain surround the cofactor and involve the α1 and α3-helices, Lα′ and Lβ1-5 loops and β1 and β2-strands (**Table S2** and **Fig. 5f**). In all cases, a hydrogen-bonding interaction between the FMN phosphate group and the side-chain of Q139 in the C-helix of CYP 1A1 was also present. NMR spectra of a membrane-anchored CYP 2B4-CPR FMN domain complex^24^ showed the same region of the FMN domain at the interface.

Site-directed mutagenesis and structural studies have been performed to identify CYP residues that interact with the redox protein for CYP 2B4 (CPR),^27,49^ CYP 17A1 (cyt b5),^50,51^ CYP 1A2 (CPR)^45^ and P450_BM3_ (FMN domain),^30^ revealing the importance of positively charged residues for redox protein binding. Even though the residues at the FMN domain binding interface of CYP 1A1 do not align completely with the interfacial residues of the other CYPs in a multiple sequence alignment (**Fig. S6**), the secondary structure regions at the interface are similar, mostly involving the B, C and L-helices, and the CC’-loop, HI-loop and the loop near the HEME. The alignment revealed that the positively charged CYP residues at the FMN domain interface were most conserved in the C3 simulation, with the conservation being highest for CYP 1A2 (80%). However, there are some differences, e.g. V267 and L270 in the H-helix of CYP 2B4 mediate CPR interactions^27^ whereas neither the corresponding aligned residues in CYP 1A1, S284 and E287, nor the H-helix make any contacts with CPR in our simulations. An NMR study of the binding of the CPR FMN domain to CYP 17A1^51^ indicated that distal residues were affected by FMN binding along with proximal residues close to those identified in our CYP 1A1 complexes (**Fig. S6**). In CYP 19A1, the B-helix (K108), C-helix (S153 and G154), K-helix (K352), K’’-L-loop (K420, Y424 and R425) and L-helix (Y441) were reported to be involved in interactions with FMN domain of CPR in a modelled CYP 19A1-CPR structure.^52^ In a model of the membrane-bound CYP2D6-CPR complex,^46^ the B-helix (R88), C-helix (R129, R133 and R140), CC’-loop (K146, K147 and S148) and loop near heme moiety (K429, E431 and R440) were found to interact with the FMN domain of CPR. In our simulations, we found that a similar, positively charged region of CYP 1A1 was involved interactions with the FMN domain of CPR (Table S2).

One of the simulations (D2’) showed complexes with a similar arrangement of the FMN and CYP domains with approximately the same protein binding faces to that observed in the crystal structure of P450_BM3_, although with the position of FMN cofactor shifted by about 20° in angle θ, see **Fig. 6a**. This rotation, together with sequence differences, permitted the cofactors to come closer to each other. Indeed, D_Fe-N5_ was much shorter in all of the CYP 1A1-CPR complexes than in the P450_BM3_ crystal structure. Recently, Dubey et al.^44^ performed MD simulations of P450_BM3_ and observed a change from a perpendicular to a parallel arrangement of the CYP C-helix and the FMN domain α1-helix. Sequence differences in the C-helix resulted in salt-bridges that prevented this reorganization in the CYP 1A1-CPR complex, **see Fig. 6b,c**. This salt-bridge interaction was observed in the C3 and D2’ simulations, whereas, in the other two simulations (C2 and D2), the FMN domain α1-helix was not in close contact with the C-helix of CYP 1A1. In the D2 simulation, the NADP domain formed transient interactions with the distal side of CYP 1A1 and in C2, the NADP domain interacted with the membrane. These two different arrangements of CPR precluded the possibility of reorganization of the C- and α1-helices as observed in P450_BM3_.

**Figure 6:**
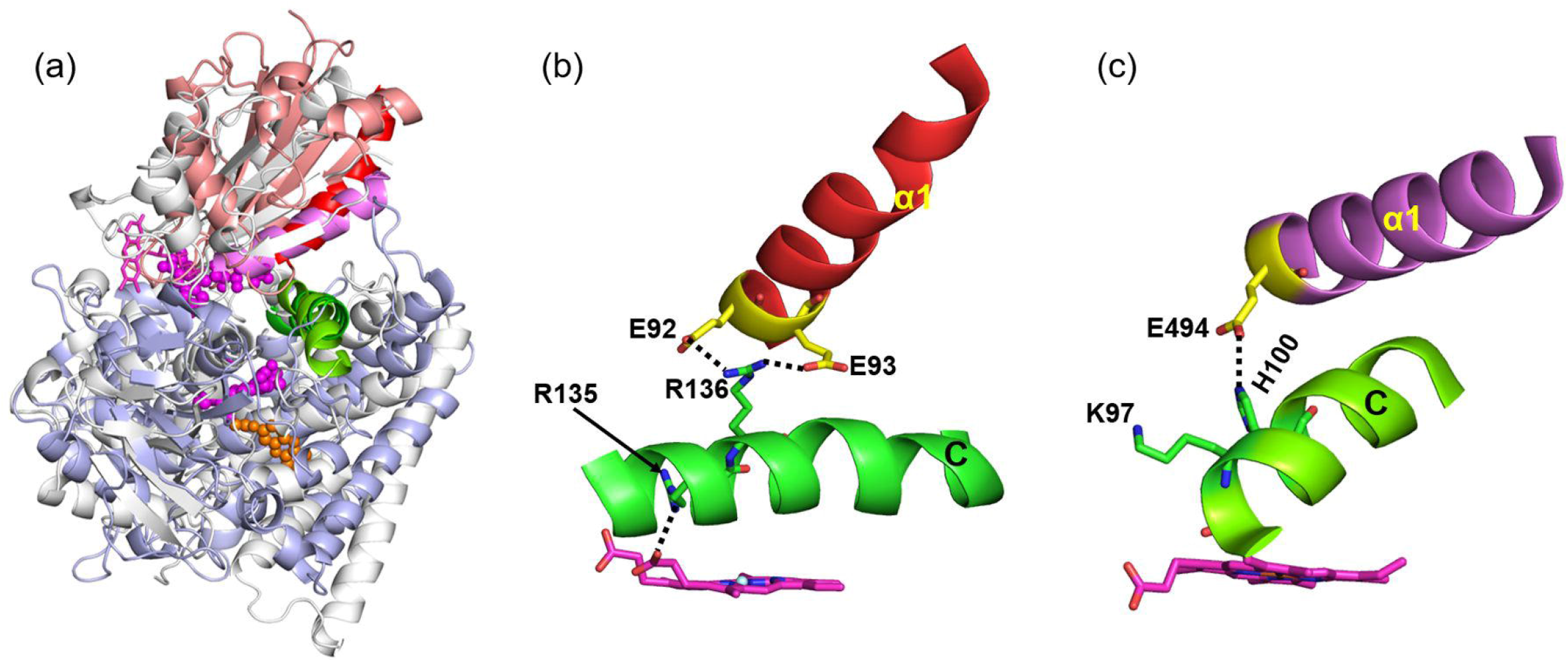
Comparison of the CYP catalytic domain-FMN domain interaction in the final CYP 1A1-CPR complex from the D2’ simulation with the crystal structure of P450_BM3_. The cofactors approach more closely in the CYP 1A1-CPR complex and the angle θ between the FMN domain α1 helix and the CYP C-helix differs by 20°. (a) Overlay of the structures by superposition of the heme cofactors. D_Fe-N5_ is 16.3 Å in the CYP1A1-CPR complex and 23.5 Å in P450_BM3_. Close-up views of (a) the simulated CYP1A1-CPR complex and (c) the crystal structure of P450_BM3_. Salt-bridges between R136 of CYP 1A1 and residues E92 and E93 of the CPR FMN domain, as well as R135 with a heme carboxylate, lock the arrangement of the FMN domain α1-helix and the CYP C-helix. In contrast, in P450_BM3_, Dubey et al.^44^ found that after MD simulation, the α1-helix reoriented and E494 lost its hydrogen-bond to H100 and approached K97 of P450_BM3_, located at the beginning of the C-helix, close enough to form a salt-bridge. This reorganization resulted in a change from a perpendicular to a parallel arrangement of the C- and α1-helices. CYP 1A1, however, lacks a corresponding positively charged residue at the N-terminus of the C-helix: P129, A132 and R135 of the C-helix of CYP 1A1 structurally align with K94, K97 and H100 of P450BM3, respectively. Color scheme: P450_BM3_ FMN and CYP domains: white; CPR FMN and CYP 1A1 globular domains : salmon and light blue, respectively; CYP C-helices: green; FMN domain α1 helices of P450_BM3_ and CPR: pink and red, respectively; FMN cofactors: pink, with P450_BM3_ and CPR cofactors in licorice and ball-and-stick representation, respectively; HEME: magenta carbons; 7-ethoxyresorufin: orange; selected residues are shown in yellow carbon representation with hydrogen bonds indicated by black dashed lines.

### Transient interactions are formed between CYP 1A1 and the NADP domain of CPR

In the initial model for the D2 system, the NADP domain of CPR was relatively far from the membrane and from the globular domain of CYP 1A1 (**Fig. 2a**). During the simulations, the NADP domain came closer to the CYP 1A1 globular domain and formed interactions (**Fig. 5c-d**). The motion of the NADP domain shifted the CPR from an open conformation towards a semi-open conformation, while retaining the center-to-center distance between the FAD and NADP domains as in the starting structure (∼34 Å). These motions of the NADP domain did not appear to have a significant impact on the interactions between CYP 1A1 and the FMN domain. There is experimental evidence supporting interactions between the NADP domain and the CYP globular domain for the CYP 1A1-CPR complex. A strong inhibition (>80%) of the CPR-supported metabolism of 7-ethoxycoumarin and ethoxyresorufin was observed in K271I and K279I single-point mutants of rat CYP 1A1, accompanied by a reduction by a factor of about 2 and 9, respectively, of the Michaelis constant for the reductase (wild-type K_m_ = 5.1 pM).^53^ Based on a combination of in vitro mutagenesis, in vivo screening and spectral analysis, it was confirmed that K268 and R275 of rat CYP 1A1 are important for CPR binding.^54,55^ However, residues 268 and 271 are in the G-helix and residues 275 and 279 are in the GH-loop, neither of which is on the proximal side of rat CYP 1A1, see **Fig. S6**.

From mutagenesis, NMR and crystallographic studies of several CYPs from several species, it is well established that the proximal side of CYP serves as an interface^25–27,49,50^ for the FMN domain. Hence, the only way for CPR to interact with the distal side of CYP 1A1 is via the NADP domain. Indeed, in the D2 simulation, the CPR NADP domain moved towards the CYP globular domain after 400 ns of simulation and residues 664, 667 and 668 formed transient contacts (with ≤ 30% occupancy) with the J-helix and the JK-loop of CYP 1A1 (**Fig. 5e**). In the D2’ simulation, the NADP domain moved towards the CYP globular domain after 200 ns and residues 664, 667 and 668 formed contacts (with ≥ 50% occupancy) with the EF-loop, the C’-helix, the G-helix and the GH-loop (**Fig. 5e**). Additionally, residues 271 and 279 of the GH-loop made transient contacts with NADP domain residues with occupancies of 9% and 13%, respectively. Thus, the interaction of the NADP domain with the G-helix and GH-loop of CYP 1A1 in the D2’ simulation is consistent with experiments.

During the simulations of the C2 complex, the FAD and NADP domains of CPR moved towards the phospholipid bilayer and started to interact with the head groups (**Fig. 5a**). Otyepka and colleagues made a similar observation in their simulations of a full-length CPR-membrane system.^56^ They found E270, P275, I307, R313 and N467 from the FAD domain and W549 and G554 from the NADP domain were points of contact with the membrane. We found the same residues making contact with the membrane in our simulations. However, the OPM^57^ webserver predicted that the FAD and NADP domains of CPR have no membrane-interacting region. The hydrophobicity and electrostatic properties of the protein surface do not indicate any membrane interaction region. Moreover, experiments indicate that the NADP domain of CPR interacts with the distal part of CYP in a way that would be incompatible with membrane binding.^7,54,55^ Hence, we discarded the C2 system from further structural analysis. The NADP domain of CPR remained further from the membrane than the CYP globular domain in all trajectories of the C3 and D2 systems.

### Simulations reveal two ET pathways and ET rates consistent with experiment

CPR transfers two electrons sequentially to CYPs during the catalytic monooxygenation cycle. The second ET step is rate-determining for most bacterial CYPs,^58^ whereas for eukaryotic CYPs, different rate-determining steps - formation of the membrane-bound CYP-CPR complex^59^, product release^36^, or the first^60^ or second ET^61^ - have been reported. The first ET to mammalian CYPs from CPR occurs at ∼2–10 s^−1 62,63^ and the second ET is usually within this range or slower.^45,62,64^

Irrespective of the different modes of interaction between CPR and CYP in the three plausible complexes (C3, D2 and D2’), only two major ET pathways were identified. These are FMN-I458-C457-HEME (56%) and FMN-K456-C457-HEME (35%), see **Fig. 5g-h**. Only the latter pathway was present in the D2’ simulation, whereas the former pathway was the predominant ET route in the other two simulations (C3: 77% and D2: 96%). Evidence supporting both these paths, which involve the backbone of the loop near the heme moiety and the heme-coordinating cysteine, comes from experiments and simulations.

As regards the first path, experiments showed that the I458V point mutation in CYP 1A1 enhanced N-demethylase activity by about two-fold due to a decreased K_m_ value, indicating slightly tighter binding of the substrate, possibly due to realignment of the heme in the mutant.^65^ The I458P mutation had a smaller and opposite effect to the I458V mutation.^65^ From a QM/MM study of CYP 3A4, it was concluded that the I443 backbone amide forms a hydrogen bond with the side chain of the heme-coordinating C442 and thus stabilizes the Fe-S bond and prevents the localization of the radical on the sulfur in the presence of substrate.^66^ Furthermore, computational modelling of CYP 19A1 and the CPR FMN domain by Magistrato and co-workers revealed that C437, A438, K440 and Y441 are involved in ET.^67^ (For alignment, see **Fig. S7**). Thus, these studies support the pathway in which the backbone of I458 in CYP 1A1 is involved in ET to the heme Fe^+2^-O_2_ species through the sulfur atom of the coordinating cysteine. With respect to the second path, the S441P mutation in CYP 17A1 (aligning with K456 in CYP 1A1) causes a complete loss of both 17α-hydroxylase and 17,20-lyase activities.^68^ Based on modelling of the membrane-bound CYP 3A4-CPR complex, Otyepka and co-workers suggested that N441, C442 and R446 of CYP 3A4 may be involved in ET.^56^ Furthermore, attachment of a photoactive Ru complex at residue 387 on the proximal side of P450_BM3_ indicated that electron transfer to the heme could occur along the peptide loop backbone from Q387 to the proximal cysteine.^69^ Superimposition of the crystal structure of P450_BM3_ on our CYP 1A1-CPR complexes indicates that Q387 is quite close to the FMN cofactor. Moreover, in the model of the CYP 2D6-CPR membrane-bound complex^46^, the loop region near the heme moiety (R440, R441, A442 and C443) was found to be involved in the electron transfer pathway, with A442 and C443 of CYP 2D6 aligning with K456 and C457, the two residues in the electron transfer pathway in our D2’ model of the CYP 1A1-CPR complex.

To investigate the relation between ET rate and the structure of the complex, 2D histograms of the computed binding free energies between the CYP globular domain and the FMN domain in the complexes (**Fig. 5i**), D_Fe-N5_ and the θ angle (**Fig. S8a-b**) were plotted against log(k_ET_). The maximum population of structures has log(k_ET_) between 0.1 to 0.7. The calculated ET rates (**Table 1**) agree well with the measured value^45^ of k_ET_ for the rat CYP 1A2-CPR complex expressed in yeast of 5.9 s^-1^. **Fig. 5i** shows that the highly populated clusters in the trajectories have intermediate binding free energies, and this is consistent with evidence from surface plasmon resonance measurements that the association of CYP and CPR in ET-compatible orientations is governed by entropic rather than enthalpic contributions.^70^ The histogram plot of D_Fe-N5_ vs log(k_ET_) (**Fig S8a**) shows that, consistent with the distances observed experimentally in several other redox proteins,^71^ D_Fe-N5_ ranges from about 13 to 17.5 Å, and that the experimental k_ET_ value is reproduced with this range of D_Fe-N5_. Notably, the histogram plot (**Fig. S8b**) of θ vs log(k_ET_) has a region from 82° to 90° where no structures were sampled during the simulations as this arrangement of the CYP 1A1 C-helix and FMN domain α1-helix is disfavored as discussed above.

### CPR affects CYP ligand tunnels due to CYP reorientation and CYP-CPR interactions

We analyzed the effects of CPR binding on the opening of ligand tunnels between the buried CYP active site and the protein surface in the conventional MD simulations described above **(Fig. 7)** and in RAMD simulations of ligand egress from the CYP active site **(Table 2)**.

**Table 2:** Relative occurrence (in %) of 7-ethoxyresorufin egress routes from the CYP 1A1 active site in RAMD simulations. Simulations were started from snapshots from the C3, D2 and D2’ simulations and the CYP 1A1-membrane system in the absence of CPR. Upon complexation with CPR, the solvent tunnel tends to close and either the 2c or the 3s tunnel tends to open.

| Tunnel | C3 | D2 | D2' | CYP<br>without<br>CPR |
| --- | --- | --- | --- | --- |
| S | 38 | 71 | 71 | 94 |
| 2ac | 0 | 4 | 4 | 0 |
| 2c | 0 | 21 | 0 | 4 |
| 3s | 58 | 0 | 25 | 0 |
| 3 | 4 | 4 | 0 | 2 |

**Figure 7:**
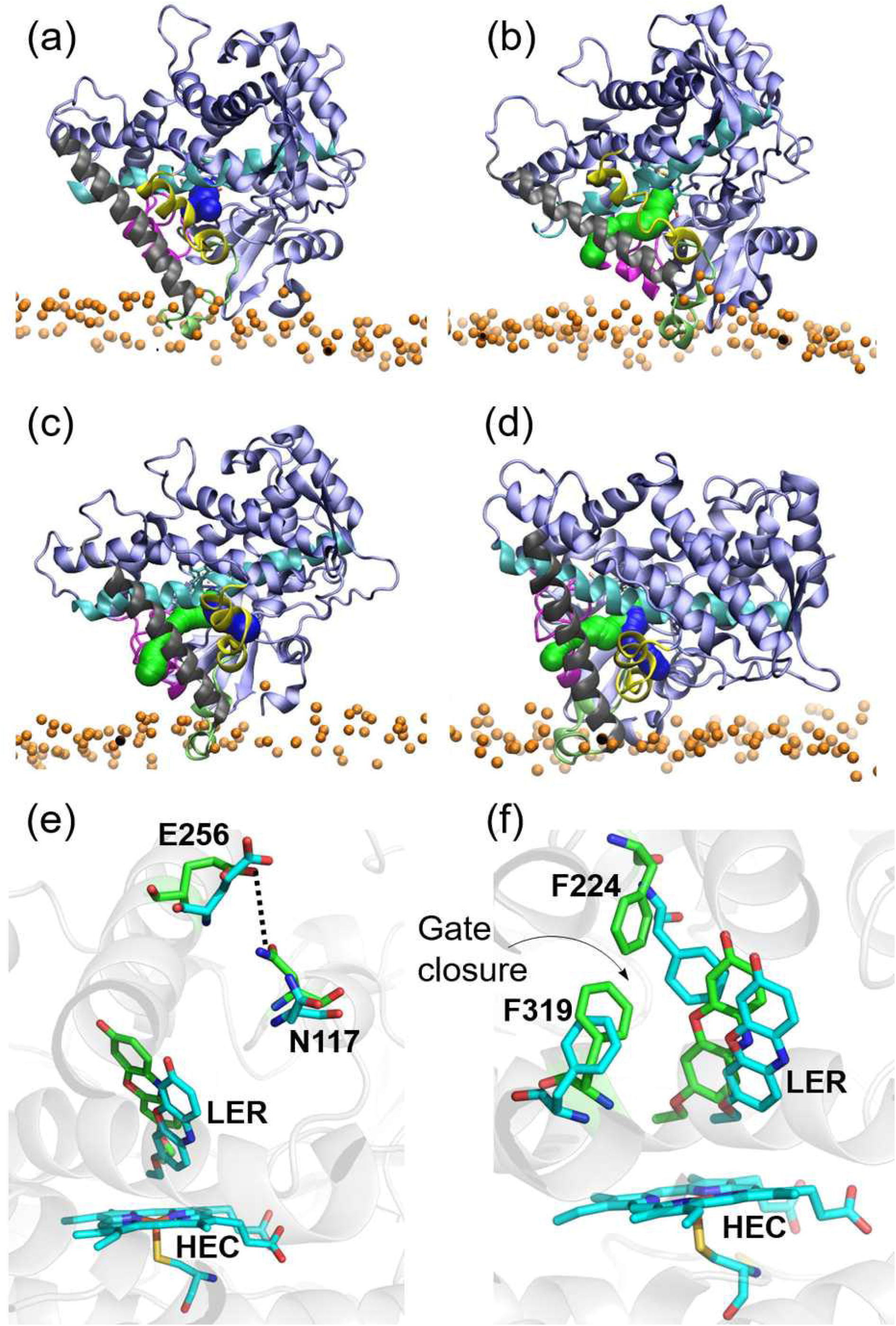
Comparison of ligand tunnels from the active site to the surface of CYP 1A1 in MD simulations of membrane-bound CYP 1A1-CPR complexes and of the CPR-free membrane-anchored-CYP 1A1. (a-d) Tunnels calculated by CAVER analysis for (a-c) three CYP-CPR simulations: (a) C3; (b) D2 (c) D2’; and (d) CYP 1A1 simulated in a membrane in the absence of CPR. (Color scheme: BC-Loop (105-128): Magenta; F-Helix (211-228): Yellow; FG-Loop (229-245): lime; G-Helix (246-272): Gray; I-Helix (304-336): Cyan. Tunnel 2ac: green; solvent tunnel: blue.) (e-f) Changes in the active site of CYP 1A1 during simulations of the CYP-CPR complexes in the membrane bilayer resulting in closure of ligand tunnels. Structures are shown before (cyan) and after (green) simulation. (e) Formation of a hydrogen bond with 95% occupancy between the side chains of N117 and E256 in the C3 simulation blocking tunnel 2ac. The occupancy of this interaction during the other simulations was less than 30%. (f) Movement of F224 and F319 in the I-helix in the D2 simulation to make van der Waals contact with an occupancy of 56%, thus blocking the route for ligand passage through the solvent tunnel. In the other simulations, the interaction between F224 and F319 was absent. Instead, there were parallel π–π interactions between the ligand and F224.

In the conventional MD simulations, two ligand tunnels in CYP 1A1 with diameters sufficient for passage of a water molecule were identified **(Fig 7a-d)**: (i) the solvent tunnel (S) between the I- and F-helices; and (ii) tunnel 2ac between the tip of the B–C loop and the G-helix (with tunnel nomenclature as in Ref ^72^). In the D2’ simulation and the simulation of CYP 1A1 in a membrane in the absence of CPR, both tunnels were open, whereas, in the C3 simulation, only the solvent tunnel was present and, in the D2 simulation, only tunnel 2ac was present. These tunnel closures were due to BC-loop motion resulting in hydrogen-bond formation closing tunnel 2ac (**Fig. 7e**) and movement of the side-chain of F224 away from the ligand to interact with F319 in the I-helix and close the solvent tunnel (**Fig. 7f**). A direct influence of CPR on these motions could not be identified by contact map analysis of the trajectories.

The RAMD simulations, in which 7-ethoxyresorufin travelled from the active site to the protein surface, revealed five ligand egress routes in the CYP-CPR complexes (S, 2ac, 2c, 3s and 3) and three routes (S, 2c and 3) in CPR-free CYP 1A1 (**Table 2)**. Although the transient interactions between F224 and F319 were perturbed in the RAMD ligand egress simulations and thus egress via the solvent tunnel was observed in all systems, the solvent tunnel tended to close and either the 2c or the 3s tunnel tended to open in the CPR-bound systems. Tunnel 2ac was rarely observed as an egress route, even though it was detected in the conventional MD simulation. Egress via tunnel 3s, between the F- and G-helices, was observed in two of the CPR-bound systems and occurred when the contacts of N221 with A250 and D253 (occupancy >90% in the CPR-free system) were broken, allowing the F- and G-helices to move apart and permit ligand passage. In summary, in these simulations, the binding of CPR to CYP 1A1 alters the distribution of egress pathways. An alteration in ligand tunnels due to CPR binding has also recently been observed in conventional MD simulations of the CYP 19A1-CPR complex in a membrane.^52^ Our results indicate, however, that more extensive studies of substrate access and product release would be necessary to confirm the effect of CPR.

## Conclusions

A multiresolution computational approach, using a combination of BD, coarse grained and all-atom MD simulations, both in the absence and presence of a phospholipid bilayer, was used for *de novo* modelling and simulation of the interactions between full-length CYP 1A1 and its redox binding partner, CPR. After sampling and selection of possible arrangements, a cumulative ∼2 μs of atomic detail MD simulations of membrane-bound CYP-CPR complexes were performed. From these simulations, we identified several arrangements of membrane-bound CPR and CYP 1A1 that are compatible with ET. Further simulations may enable a complete sampling of the configurational ensemble of the membrane-bound CYP 1A1-CPR complex but already, from the simulations done, we have been able to identify two main interaction modes consistent with available mutagenesis data and having ET pathways compatible with experimentally observed ET rates.

Upon binding CPR, the interaction between CYP 1A1 and the membrane became weaker as the CYP-FMN domain interface area increased. Although no TM-helix-TM-helix contacts between CYP and CPR were observed, the large interface area of the final complexes (ca. 2000 Å^2^) was consistent with the high binding affinity of the CYP 1A1-CPR complex. In all the final model complexes, the CYP1A1 catalytic domain reoriented in the membrane. This reorientation affected ligand access and egress pathways between the active site and the membrane but there was little effect on interactions of the ligand in the catalytic site on the timescale of the simulations. CPR rearranged in the membrane and underwent a large conformational change from the open to the semi-open form which enabled formation of an additional interface between the distal side of CYP 1A1 and the CPR NADP domain. These data clearly indicate that, in presence of CPR, a rearrangement in orientation of CYP occurs and vice versa.

The FMN domain binding face composition of CYP1A1 in the modelled CYP 1A1-CPR complexes is mostly similar to other experimentally determined FMN domain binding interfaces of CYPs.^27,30,45,49,51^ One of our model complexes (D2’) has a similar arrangement of the CPR FMN domain with CYP 1A1 to that observed in the crystal structure of P450_BM3_, although, with the position of the FMN cofactor shifted by about 20°. The computed ET rates and pathways from CPR to CYP 1A1 are in excellent agreement with available experimental data. The slower ET rates compared to soluble CYPs appear to result from the competing requirements of ET and membrane binding by both CYP and CPR, as well as sequence differences at the CYP-FMN domain interface. The importance of identifying ET pathways is highlighted by the fact that mutation of I458, a key residue in the dominant computed ET pathway, has been shown to affect prodrug activation.^65^ The most common non-synonymous polymorphisms of CYP 1A1 have the mutations T461N and/or I462V, resulting in high-risk variants for non-small cell lung cancer in non-smokers.^73,74^ We observed direct contact (<5 Å) of I462 with I458 only in our simulations in the presence of CPR, indicating that the I462V variant may have a role in the regulation of substrate metabolism. T461 interacted with the Lβ4-loop region of the CPR FMN domain in our simulations, indicating that alterations in CPR binding of the T461N mutant might contribute to the observed increased formation of the mutagenic diol epoxide of benzo[a]pyrene-7,8-dihydrodiol^75^. Thus, the present simulations give detailed atomistic insights into structural and functional aspects of CYP 1A1-mediated drug and carcinogen metabolism. Moreover, they provide insights into how CYP-redox partner interactions can vary, and provide a basis for further work to explore the interactions of different CYPs with CPR.

## Methods

The computational workflow for building systems for simulations of full-length CYP-CPR complexes in a membrane bilayer is illustrated in **Fig. 1** and described briefly below; further details are given in **Supplementary Methods** and **Fig. S1**.

### Protein structures

Modeling of CYP 1A1 was based on the crystal structure of the globular domain of human CYP 1A1 (PDB ID: 4I8V; 2.6 Å resolution), in complex with the inhibitor α-naphthoflavone,^76^ following refinement and rebuilding using the PDB-REDO server^77,78^ and addition of missing residues using Modeller^79–81^ v.10. All four chains (A-D) of the asymmetric unit were used in BD simulations. For the full protein, the TM-helix in CYP 1A1 was assigned to residues 6-27 and the flexible linker and TM-helix were modelled as described previously. ^57,82^The structure of an open conformation of human CPR was modelled on the basis of the crystal structures of the N-terminally truncated rat CPR in an open conformation (PDB ID: 3ES9 chain B; 3.4 Å resolution)^62^ and the closed form of the N-terminally truncated human CPR (PDB ID: 3QE2; 1.75 Å resolution).^83^ Missing residues were modelled using VMD^84^ and Modeller.

### Brownian dynamics simulations

BD rigid-body docking simulations of the four structures of the globular domain of CYP 1A1 and the N-terminally truncated CPR were carried out using the SDA 7 software.^39,85–87^ All cofactors were retained but the ligand was removed from the CYP. Docking was performed subject to electrostatic interaction, electrostatic desolvation, and non-polar desolvation forces.^85–87^

240,000 independent trajectories were run starting with CPR at a random relative orientation and position at a center-to-center distance of 450 Å from the CYP, and they were all terminated when the inter-protein center-to-center separation reached 600 Å. ET in redox proteins typically occurs when the edges of the redox centers lie within 14–15 Å.^71^ Hence, the coordinates of diffusional encounter complexes were recorded subject to two distance constraints: (i) the cofactor-cofactor distance from FMN:N to HEME:Fe, D_Fe-N5_ < 20 Å, and (ii) the center-to-center distance between the CYP globular domain and the CPR FMN domain, D_CYP-FMN domain_ < 60 Å, to ensure close approach of the domains.

For each of the four CYP structures, the energetically ranked top 5,000 docked encounter complexes were clustered into 10 clusters ranked by cluster size using a hierarchical method.^39^ Thus, 40 representative encounter complexes were generated and named according to the chain identifier in the crystal structure of CYP 1A1 and the docking cluster number of CPR. E.g., C3 for chain C of CYP 1A1 and the third largest docked cluster of CPR. These complexes were further filtered according to interaction energy, and to ensure that the clusters were structurally distinct and did not have orientations of CPR inconsistent with membrane anchoring if the CYP was assumed to be positioned in the membrane as observed in MD simulations of CYP 1A1 in a POPC bilayer. As a result, six structures of diffusional encounter complexes were selected for further studies.

### All-atom molecular dynamics simulations in aqueous solution

To account for protein flexibility, all-atom MD simulations in 150 mM aqueous solution (referred to as “soluble simulations”) were carried out for all 6 representative structures of CYP 1A1 □ CPR encounter complexes obtained from BD simulations. For computational efficiency, only the globular domain of CYP 1A1 and the FMN domain of CPR were simulated in the soluble simulations. After equilibration, all-atom MD production simulations were performed for about 50 to 140 ns using the NAMD v2.10 software.^88^

### All-atom molecular dynamics simulations of membrane-bound full-length CYP-CPR systems

All the simulations with phospholipid bilayers, “membrane simulations”, were carried out for complete structures of full-length CYP and CPR. The initial configurations of the complexes of the globular domain of CYP and the FMN domain of CPR were obtained from the last frames of the soluble simulations.

#### a) Preparation of full-length CYP structures

We previously developed a protocol for inserting the full-length structure of a CYP into a membrane bilayer without predetermining the orientation or insertion depth.^16^ Hence, we superimposed the CYP domain of the CYP_globular-domain_ □ CPR_FMN-domain_ complexes onto a modelled structure of the membrane-bound full-length CYP 1A1 in a POPC membrane.^82^ Next, we replaced the globular domain of CYP 1A1 with each CYP_globular-domain_ □ CPR_FMN-domain_ complex and connected the TM-helix and the linker region (residues 1 to 50) from the membrane-bound CYP 1A1 simulated in the absence of CPR with residue H51 of CYP 1A1 in the CYP_globular-domain_-CPR_FMN-domain_ complex.^89^ A local structural refinement to optimize the covalent linkage between residues 50 and 51 was performed using the interactive energy minimization module of Maestro (Schrödinger, LLC, New York, NY, 2015)^89^, resulting in complete initial structures of the membrane-bound CYP_full-length_ LJ CPR_FMN-domain_ complex.

#### b) Preparation of full-length CPR structures

The FMN domain of CPR (residues 66 to 230) in each membrane-bound CYP_full-length_ LJ CPR_FMN-domain_ complex was superimposed onto the FMN domain of truncated human CPR (residues 66 to 677) and residues 231-677 of CPR were extracted and connected with the FMN domain of the membrane-bound CYP_full-length_ LJ CPR_FMN-domain_ complex. The amide bond between residues 230 and 231 was optimized using Maestro. The TM-helix region of CPR was predicted by using the Orientation of Protein in Membrane (OPM) (http://opm.phar.umich.edu/server.php)^57^ and the TMpred^90^ webservers and its structure was modelled using Modeller v 9.10. Next, the modelled TM-helix (residues 26 to 50, with residues 26 to 44 being helical) was placed arbitrarily in the membrane so that there were no interactions between the CYP and CPR transmembrane helices. Then, the missing residues (1-25 of the membrane binding domain and 51-65 of the flexible tether region) were added using Modeller to complete the full structure of the membrane-bound CYP_full-length_ □ CPR_full-length_ complex.

#### c) Simulation setup and trajectory generation

Each CYP-CPR complex in a bilayer of POPC phospholipids was immersed in a periodic box of water molecules with 150 mM ionic strength maintained by adding Na^+^ and Cl^−^ ions. Production simulations were performed for protein-phospholipid bilayer systems based on three encounter complexes (with one system simulated in duplicate) using the NAMD v2.10 software.

### Post-analysis of MD trajectories

CYP globular domain-FMN domain interaction free energies were estimated using the MMGBSA^91,92^ protocol implemented in the AMBER 14 package (without performing normal mode analysis for estimating the vibrational entropy contribution) for snapshots collected at 100 ps intervals after completion of 350 ns simulation (i.e. for the last 150-200 ns of the trajectories of the protein complexes). Substrate access tunnels were computed using CAVER 3.0^93^ for the last 500 snapshots of each trajectory collected at 100 ps intervals. Random acceleration molecular dynamics (RAMD)^94^ simulations, as implemented in NAMD 2.10, were performed to study the ligand egress pathways from the CYP 1A1 active site in the presence and absence of CPR. ET pathways and rates were computed using Beratan’s model with the VMD pathway plugin module.^95^

## Supporting information

MSExcel sheet of encounter complexes

Data for Figs 3,4 and 5i and Tab. 1

## AUTHOR CONTRIBUTIONS

GM, PPN and RCW designed the research; GM performed computations and GM, PPN and RCW analyzed the computational data; GM and RCW wrote the manuscript.

### ACKNOWLEDGEMENTS

We thank the Klaus Tschira Foundation, the German Academic Exchange Service (DAAD) (scholarship to PPN), and the Center for Modelling and Simulation in the Biosciences (BIOMS) (postdoctoral fellowship to GM) for support. The authors acknowledge support for computing resources from the state of Baden-Württemberg through bwHPC and the German Research Foundation (DFG) through grant INST 35/1134-1 FUGG and for use of the Hazel Hen Cray XC40 at the high-performance computing center, Stuttgart, Germany (HLRS; Project Dynathor). The NAMD software was developed by the Theoretical and Computational Biophysics Group in the Beckman Institute for Advanced Science and Technology at the University of Illinois at Urbana-Champaign. We thank Dr. Stefan Richter for assistance in optimizing software performance, Dr. Xiaofeng Yu for preliminary simulations of CYP-CPR systems, Dr. Ghulam Mustafa for help in setting up CYP simulations, and Soren von Bülow for implementing scripts for computing inter-protein ET pathways.

## Supporting Information Available

Supplementary Methods, Figures S1-S10, Tables S1-S4, MS Excel sheet listing encounter complexes and file (rar) containing data for figures 3, 4 and 5i and Table 1.

## Data Availability

Atomic coordinates from the last frames of the simulations of three CYP-CPR complexes are available at ModelArchive: C3: doi:10.5452/ma-3oc22, D2: doi:10.5452/ma-8dk69 and D2’: doi:10.5452/ma-7au7g. The datasets generated during and analyzed during the current study are available from the corresponding author on reasonable request.

