## Supplementary Information for "An electron transfer competent structural ensemble of membrane-bound cytochrome P450 1A1 and cytochrome P450 oxidoreductase"

**Supplementary Methods** ........................................................................................ **1-24**

- Details of the protein structure preparation………………………………………….... **1**
- Details of Brownian dynamics (BD) simulation protocol .............................................. **1-3**
- Details of molecular dynamics (MD) simulation protocols............................................ **3-6**
  - *Protocol for MD simulations of CYP-FMN domain complexes in aqueous solution (‘soluble’ simulations)* ......................................................................... 3-4
  - *Protocol for MD simulations of CYP-CPR complexes in a membrane bilayer* ... 4-5
  - *Protocol for Random Acceleration Molecular Dynamics (RAMD) simulations* . 5-6
- Electron Transfer (ET) pathway and kinetics calculation............................................... **6-7**

**Supplementary Figures** ........................................................................................ **8-14**

**Supplementary Tables** ........................................................................................ **15-20**

**Supplementary References**.................................................................................... **21-24**

### Supplementary Methods

#### *Details of protein structure preparation*

*Preparation of the open form of human CPR*

The FMN domain and, separately, the FAD and NADP domains of the crystal structure of the closed form of the N-terminally truncated human CPR (PDB ID: 3QE2; 1.75 Å resolution),[1] were superimposed onto chain B of rat CPR. The hinge region (residues 231-243) of human CPR was also superimposed onto the hinge region of the crystal structure and the missing residues (236-239) were then modeled in VMD.[2] Thus, we obtained a complete structure of the TM-helix-truncated human CPR in an open form. However, the sequence of the TM-helix-truncated human CPR in the crystal structure (PDB ID: 3QE2) is only 99% identical to the sequence reported in Uniprot (ID: P16435). Hence, we performed homology modelling using Modeller v.10 with the aforementioned structure as a template to model the human TM-helix-truncated CPR with the aforementioned Uniprot sequence.

#### *Details of the Brownian dynamics (BD) simulation protocol*

Translational and rotational diffusion coefficients were computed for each protein using HYDROPRO.[3] The force field parameters for the CPR cofactors were assigned with the generalized Amber force field (GAFF)[4] and the parameters of the heme were assigned as described previously.[5,6] The partial atomic charges of the cofactors were derived by the RESP fit method using the R.E.D. server.[7] PDB2PQR[8,9] was used to assign hydrogen atoms, van der Waals radii and partial atomic charges at pH 7 to the structures of CYP 1A1 and CPR according to the AMBER force field,[4,10] resulting in a total net charges for CYP 1A1 (residues 36-512 and HEME) of +3e, and CPR (residues 66-677 and FMN, FAD and NADP) of -26e. Then, the molecular electrostatic potentials of the proteins were computed by solving the linearized Poisson-Boltzmann equation using the APBS[11] software on cubic grids extending 161 Å for the CYP and 193 Å for the CPR with a grid spacing of 1 Å. The dielectric boundary was defined by the protein's van der Waals surface using the parameters smol, srad = 0.0 and swin = 0.3, with the protein interior assigned a dielectric constant of 2.0 and the solvent assigned a dielectric constant of 78.0. An ionic strength of 150 mM was assigned with a temperature of 298.15 K with an ion radius of 1.5 Å. Effective charges on each protein were calculated using the ECM module of SDA 7.[12–15] Effective charge sites were located on the charged residues and the polar atoms (O, N, S, Fe) of the cofactors. Electrostatic and non-polar desolvation grids of the same size as the electrostatic potential grids were generated with a grid spacing of 1 Å, and an ionic strength of 150 mM was assigned to compute the electrostatic desolvation grid. Docking was performed using the interaction grid multiplication factors 0.5, 0.36 and -0.013 for the electrostatic interaction, electrostatic desolvation, and non-polar desolvation terms,[12–14] respectively. Similar results were obtained in calculations with the same parameters but using an electrostatic desolvation term with a grid multiplication factor of 1.67.[14] To avoid overlap of the proteins during the simulations, an exclusion grid was computed for each protein with a grid spacing of 0.25 Å and a probe radius of 1.77 Å.

A total of 240,000 independent trajectories, 60,000 for each of the 4 chains of the CYP 1A1 crystal structure, were generated. All trajectories were started with CPR at a random relative orientation and position at a center-to-center distance of 450 Å from the CYP (which was held stationary), and they were all terminated when the inter-protein center-to-center separation reached 600 Å. A variable timestep was used with minimum and maximum time steps of 0.5 and 20 ps, respectively. During BD docking simulations, two distance constraints were imposed for recording the coordinates of diffusional encounter complexes: (i) the cofactor-cofactor distance, i.e.; from N of FMN in CPR to Fe of HEME in CYP, $D_{Fe\text{-}N5}$ < 20 Å, and (ii) the center-to-center distance between the globular domain of CYP and the FMN domain of CPR, $D_{CYP\text{-}FMN\ domain}$ < 60 Å, , to ensure close approach of the domains..

The energetically ranked top 5,000 docked encounter complexes differing by more than 1 Å CPR root mean squared deviation (RMSD) from each other that satisfied the aforementioned constraints were recorded, along with their populations (number of times sampled), during the simulations. Subsequently, the structures were clustered into 10 clusters ranked by cluster size using a hierarchical method.[15] Thus, a total of 40 representative encounter complexes (1 for each of the 10 clusters for each of the 4 chains) was generated for the CYP 1A1-CPR system. The complexes were named according to the chain identifier in the crystal structure of CYP 1A1 and the docking cluster number of CPR. E.g., an encounter complex between chain C of CYP 1A1 and the representative structure of the third largest docked cluster of CPR was named C3.

These complexes were further filtered according to interaction energy and the top 15 complexes with a cluster size (CLsize) exceeding 100 were selected. As the orientation of the FMN domain of CPR with respect to CYP 1A1 was similar among some of these complexes, a further criterion was added, namely that if the RMSD of the FMN cofactor atoms between two different encounter complexes, $RMSD_{FMN}$, was less than 4 Å, the complex with the lowest distance, $D_{Fe-N5}$, was selected. After applying these two criteria, 7 complexes were removed. A further two complexes were removed because they had orientations of CPR inconsistent with membrane anchoring if the CYP was assumed to be positioned in the membrane as observed in MD simulations of CYP 1A1 in a POPC bilayer (**Fig. S2**): either the distance of the N-terminus of the FMN domain to the membrane surface was too long to be connected by the flexible linker (21 residues) to the CPR TM-helix (73.3 Å in the A6 complex) or the FAD or NADP domain approached too close to the membrane surface (within 12 Å of a lipid phosphorous atom) (B2 complex in which the FAD domain was within 11 Å of the membrane surface). Finally, six structures of diffusional encounter complexes for the CYP 1A1-CPR system were selected for further studies.

#### *Details of molecular dynamics (MD) simulation protocols*

*(a) Protocol for MD simulations of CYP-FMN domain complexes in aqueous solution ('soluble' simulations).*

The AMBER ff14SB force field[10] was used for the protein. The bonded parameters for all cofactors and substrates were obtained from GAFF[4] and the literature.[5,6] An ionic strength of 150 mM was obtained by adding $Na^+$ and $Cl^-$ ions to a rectangular periodic box of TIP3P[16] water molecules. 16 excess $Na^+$ ions were added to neutralize the systems with charges of +3e for the CYP 1A1 globular domain and -19e for the CPR FMN domain.

All the systems were energy minimized using the AMBER v14 software.[17,18] MD equilibration and production runs were performed using the NAMD v2.10 software[19] starting with the AMBER minimized coordinates. The energy minimization of the entire system (the complex of the CYP 1A1 globular domain (residues 36 to 512) and the FMN domain of CPR (residues 66 to 230) a box of water molecules and ions) was carried out by restraining non-hydrogen protein atoms with a force constant that gradually decreased from 1000 to 0 kcal/mol·Å$^2$ for each system. During energy minimization, the maximum number of cycles was set to 14000 steps: 1400 steps steepest descent followed by 12600 steps conjugate gradient. Minimization was stopped when the root mean square energy gradient for the input coordinates was less than 0.0001 kcal/mol/Å. The system was then equilibrated at constant pressure and temperature (NPT) for a total of 12.8 ns with a time step of 1 fs for the first 7.8 ns and a time step of 2 fs for the remaining 5 ns at 310 K using the Particle Mesh Ewald (PME) method for long range electrostatic interactions with a non-bonded cut-off of 10 Å. The SHAKE algorithm[20] was imposed to constrain all the bonds to hydrogen atoms. The Langevin dynamics method with a damping coefficient of 5/ps for the first 2.8 ns and 1/ps for the remaining time was used for temperature control. The Nosé-Hoover Langevin piston was used for pressure control with an oscillation time of 100 fs and a damping time of 50 fs for the first 2.8 ns and 500 fs for the rest of the equilibration. The first 1.8 ns of equilibration was carried out by gradually reducing the restraint force constant from 100 to 0.01 kcal/mol·$Å^2$ on all non-hydrogen atoms of the protein residues. Then the remaining 11 ns of equilibration were run without harmonic restraints. The subsequent production runs were performed for about 50 to 140 ns, depending on the convergence of structural parameters, with a time step of 2 fs and a temperature of 310 K in an NPT ensemble. Structural analysis was carried out for the soluble simulations using AMBER CPPTRAJ[21] for snapshots collected at 2 ps intervals. Xmgrace (plasma-gate.weizmann.ac.il/Grace/) was used for plotting. Molecular visualization was performed with PyMOL (https://pymol.org/2/) or VMD.

*(b) Protocol for MD simulations of CYP-CPR complexes in a membrane bilayer*

The initial coordinates of the membrane were taken from the simulated equilibrium geometry of the membrane-bound CYP 1A1 in the absence of CPR. Therefore, to accommodate the CPR, the membrane size was increased by 21 Å and 43 Å along the y-axis for the C2 and C3 systems, respectively, and by 29 Å along the x-axis for the D2 system. POPC phospholipids were assigned AMBER Lipid14 force field parameters.[22]

All the systems were energy minimized using the AMBER v14 software. MD equilibration and production runs were performed using the NAMD v2.10 software with the AMBER energy minimized coordinates. The energy minimization of the entire system ($CYP_{full\text{-}length}$– $CPR_{full\text{-}length}$ complex) was carried out by restraining non-hydrogen protein atoms with a harmonic force constant that gradually decreased from 1000 to 0 kcal/mol·$Å^2$. During energy minimization, the maximum number of cycles was set to 180,000 steps : 133,000 steps steepest descent followed by 47,000 steps conjugate gradients, and minimization was stopped when the the root mean squared energy gradient for the input coordinates was less than 0.0001 kcal/mol/Å. The system was then equilibrated at constant pressure, temperature and surface area (NPAT) for a total of 11.4 ns with a time step of 0.5, 1 and 2 fs for 0 to 1.4, 1.4 to 6.4, and 6.4 to 11.4 ns, respectively, at 310 K using the Particle Mesh Ewald (PME) method for long-range electrostatic interactions with a non-bonded cut-off of 10 Å. The SHAKE algorithm was imposed to constrain all the bonds to hydrogen atoms. The Langevin dynamics method with a damping coefficient 5/ps for the first 1.4 ns and 1/ps for the rest of the equilibration was used for temperature control and the Nosé-Hoover Langevin piston was used for pressure control with an oscillation time of 100 fs and a damping time of 50 fs for the first 1.4 ns and 500 fs for the rest of the equilibration. Initially, a 900 ps equilibration run was carried out by gradually reducing the restraint force constant from 100 to 0.01 kcal/mol·Å$^2$ on all non-hydrogen atoms of the protein residues. Then, the remaining 10.5 ns of equilibration were run without harmonic restraints. Subsequent production MD runs were performed at constant pressure and temperature (NPT ensemble) for around 500 ns while maintaining a constant ratio of bilayer x-y dimensions to ensure that the protein complex was surrounded by sufficient lipid in all directions in the plane of the membrane. Structural analysis was carried out for the membrane simulations using VMD scripts for snapshots collected at 20 ps intervals. The ParmED tool (https://github.com/ParmEd/ParmEd) was used to convert the NAMD trajectories to Gromacs compatible formats for contact map analysis with the CONAN[23] tool. All other aspects of the protocol for the production runs were the same as for the soluble simulations.

#### *(c) Protocol for Random Acceleration Molecular Dynamics (RAMD) simulations*

The RAMD simulations were carried out with an additional randomly oriented acceleration of 0.035 kcal/Å.g applied to the center of mass of 7-ethoxyresorufin in the membrane-bound CYP 1A1-CPR complex. The maximum run time of each RAMD simulation was 2 ns. The same MD simulation parameters as given above were used for the RAMD simulations. The distance travelled by the ligand was monitored at 100 fs (50 steps with a timestep of 2 fs) intervals. If the distance travelled was less than a pre-defined minimum distance of 0.025 Å, the direction of the force was changed to another randomly chosen direction. When the total distance travelled by the ligand was more than 40 Å, the RAMD simulation was terminated and the ligand was considered to be completely outside the CYP. For each model studied, the last snapshot from the production run of the membrane-bound CYP 1A1-CPR complex was used as the initial structure for performing RAMD simulations. 24 independent RAMD simulations were run for each of the three membrane-bound CYP 1A1-CPR complexes and for membrane-bound CYP 1A1 in the absence of CPR. Results were analyzed based on visualization of RAMD trajectories by VMD.

***Electron Transfer (ET) pathway and kinetics calculation***

Beratan's model as implemented in the VMD pathway plugin module was used to compute electron tunneling pathways and ET rates (https://people.chem.duke.edu/~ilya/Software/Pathways/docs/pathways.html).[24] This model identifies an effective ET tunneling pathway between two redox centers (here, N5 of FMN to Fe of heme) by evaluating the highest donor-to-acceptor coupling constant ($T_{DA}$) value. In this model, it is considered that the electron can migrate from the donor to the acceptor center through a covalent bond, a hydrogen bond or vacuum, with different migration probability penalties. The final value of $T_{DA}$ is obtained as the product of the penalties for all of the steps involved. The concept of electronic coupling was given by Marcus in his theory[25–27] for estimating the rate of outer sphere ET. Another two prerequisite parameters in the Marcus model are the driving force of the reaction, $\Delta G^0$, and the reorganization energy of the reactants, $\lambda$. The value of $\Delta G^0$ can be calculated from the redox midpoint potential differences of the donor (FMN in CPR) and acceptor (Heme in CYP) in the CYP-CPR complex.

In order to compute the ET rate, we considered the second ET to be the rate-limiting step in the catalytic cycle of CYPs (eq. 1):,

$(O_2)Fe^{2+}$-Porphyrin + $FMN_{sq} \rightarrow (O_2^{1-})Fe^{2+}$-Porphyrin + $FMN_{ox}$ ......(eq 1)

We therefore estimated the redox potential values for $FMN_{ox/sq}$ (sq and ox are the semiquinone and oxidized forms of FMN, respectively) and $(O_{2(-1/0)})Fe^{2+}$-Porphyrin (HEME-oxo). The redox potential value for $FMN_{ox/sq}$ of human CPR in a POPC membrane has not been reported. However, the corresponding value for rat CPR in a POPC membrane has been reported as -0.016 eV.[28] Owing to the very high sequence similarity (92%), the redox potential value is expected to be very similar for human CPR in a POPC bilayer. We, therefore, adopted this value of -0.016 eV for the redox potential of $FMN_{ox/sq}$ in human CPR. The redox potential value of the HEME-oxo species is not available in the literature for any isoforms of CYP. However, experimental evidence showed that cytochrome b5 (cyt b5) is capable of transferring the second electron to the heme of CYP, but not the first electron in the catalytic cycle. The redox potential value of human cyt b5 is within the range from 0.0205[29] to 0.025[30] eV vs NHE as measured by experiment. Therefore, the redox potential value of the HEME-oxo moiety in any isoform of CYP is expected to be at least the same or more positive than the value for cyt b5 in order to have a favorable ET reaction. Therefore, we considered the redox potential value for HEME-oxo in CYP 1A1 to be 0.025 eV. Thus, the driving force for the second stage of electron transfer for the CYP 1A1-CPR complex was assigned as -0.041 eV. The reported values of the reorganization energy (λ) for $P450_{BM3}$, typically range between 1 to 1.2 eV.[31] For the present study, we have therefore considered λ to have a value of 1.2 eV.

ET rates were computed for the last snapshot of the soluble simulation trajectories. Average ET rates and the distribution of ET pathways were computed for snapshots collected at 100 ps intervals for the last 350 ns of each of the membrane simulations. A total of 6272 snapshots from the simulations of the C3, D2 and D2' systems were also analysed to compute the overall percentage use of each ET pathway.

**Supplementary Figures**

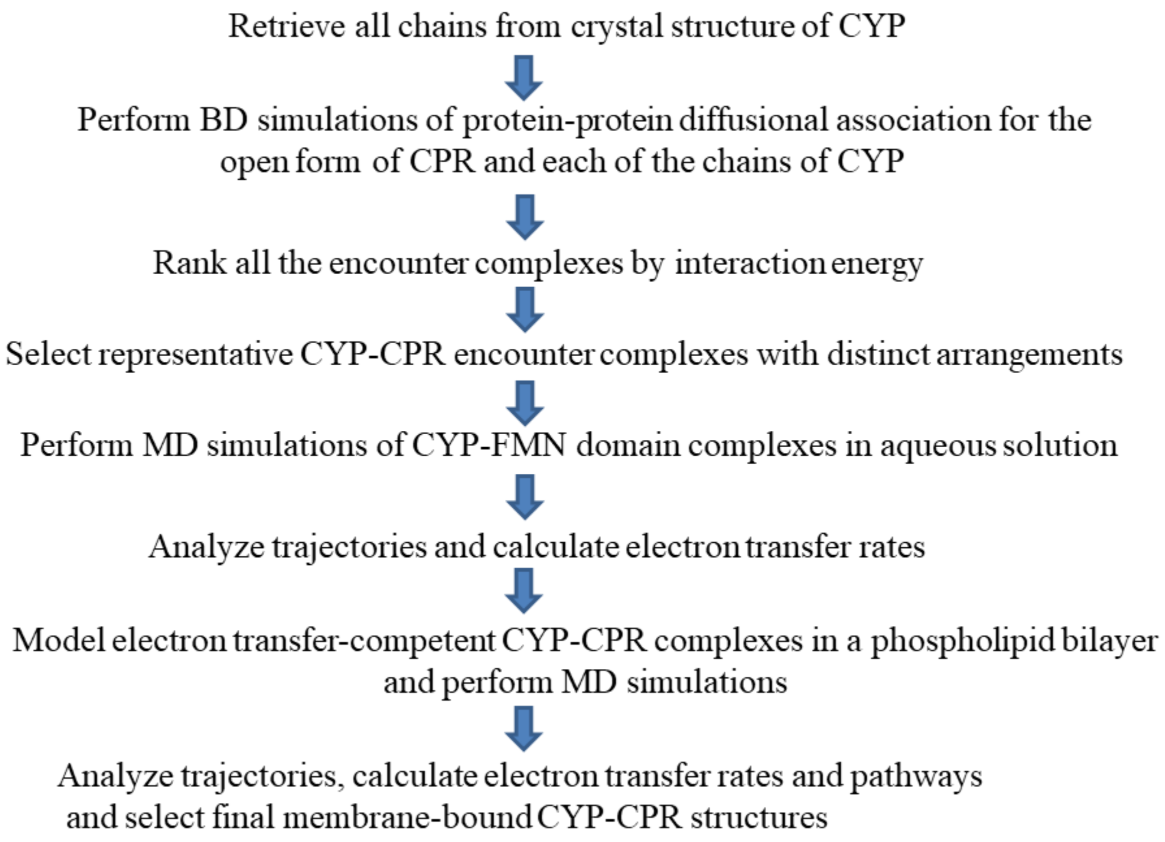


**Figure S1:** Computational workflow for modeling and simulating a CYP-CPR complex in a phospholipid bilayer.

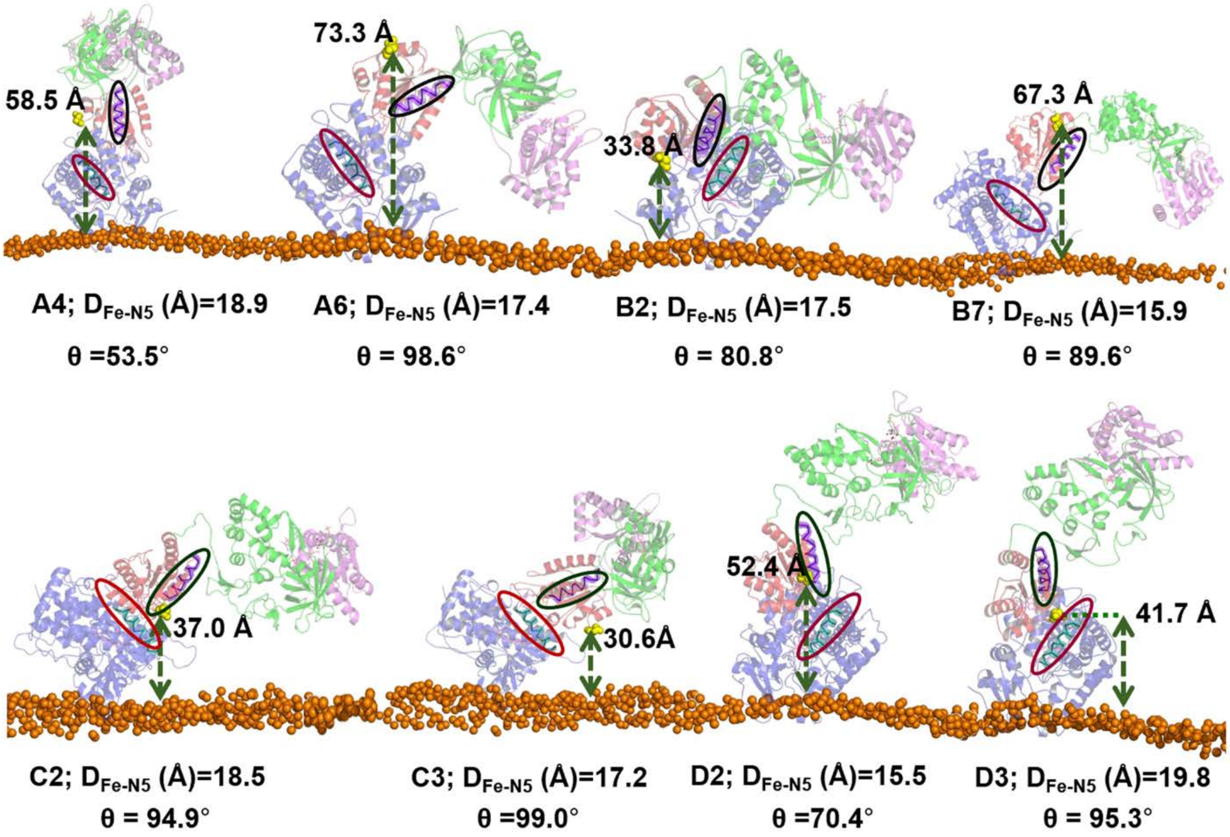


**Figure S2:** Orientations with respect to the membrane of eight CYP 1A1-CPR encounter complexes obtained from the rigid-body BD docking simulations between the globular domains of CYP 1A1 and CPR. Each encounter complex is named according to crystal structure chain identifier of the CYP 1A1 structure docked (A-D) and the cluster number that it represents (ClX, where X is 1-10). These eight clusters were obtained after filtering the 40 docked clusters according to interaction energy and to ensure that the clusters were structurally distinct. The orientations were obtained by superimposing the CYPs of the BD-docked complexes onto the membrane-bound CYP 1A1 obtained from MD simulations. The proteins are shown in cartoon representation (CYP 1A1: blue; FMN domain: red; FAD domain: green; NAD domain: magenta) and the cofactors in magenta stick representation. Phosphorus atoms in the head groups of the upper phospholipid monolayer are shown by orange spheres. $D_{Fe\text{-}N5}$ is the redox center separation distance in Å. The green arrows indicate the distance (Å) of the N-terminus of the FMN domain from the bilayer surface. Helices C of CYP 1A1 and α1 of the FMN domain are highlighted by red and black ellipses, respectively, and θ is the angle between the two helices. Six encounter complexes were selected for molecular dynamics simulations. Two clusters were discarded as they had orientations of CPR inconsistent with membrane anchoring if the CYP was assumed to be positioned in the membrane as observed in MD simulations of CYP 1A1 in a bilayer: A6 due to the large distance from the N-terminus of the FMN domain to the bilayer that would need to be spanned by the CPR flexible linker, and B2 due to the proximity of the FAD and NADP domains to the membrane (atoms within 11 Å).

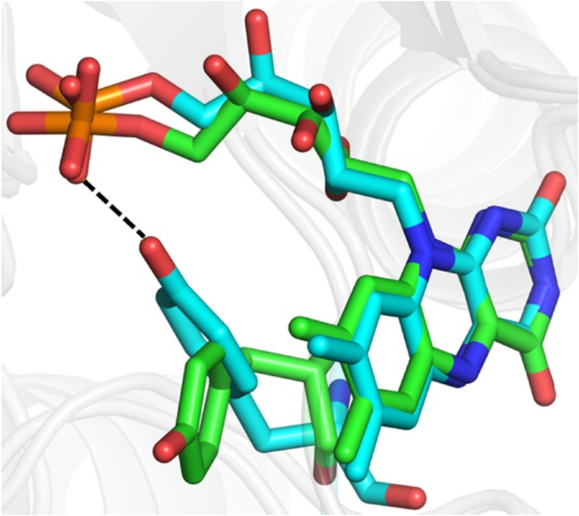

**Figure S3:** Position of Y140 of CPR and the FMN cofactor before (cyan) and after (green) MD simulation of the A4 complex. The hydrogen bond between the side chain of Y140 and the phosphate group of the FMN cofactor is shown by a yellow dashed line and is lost during the simulation.

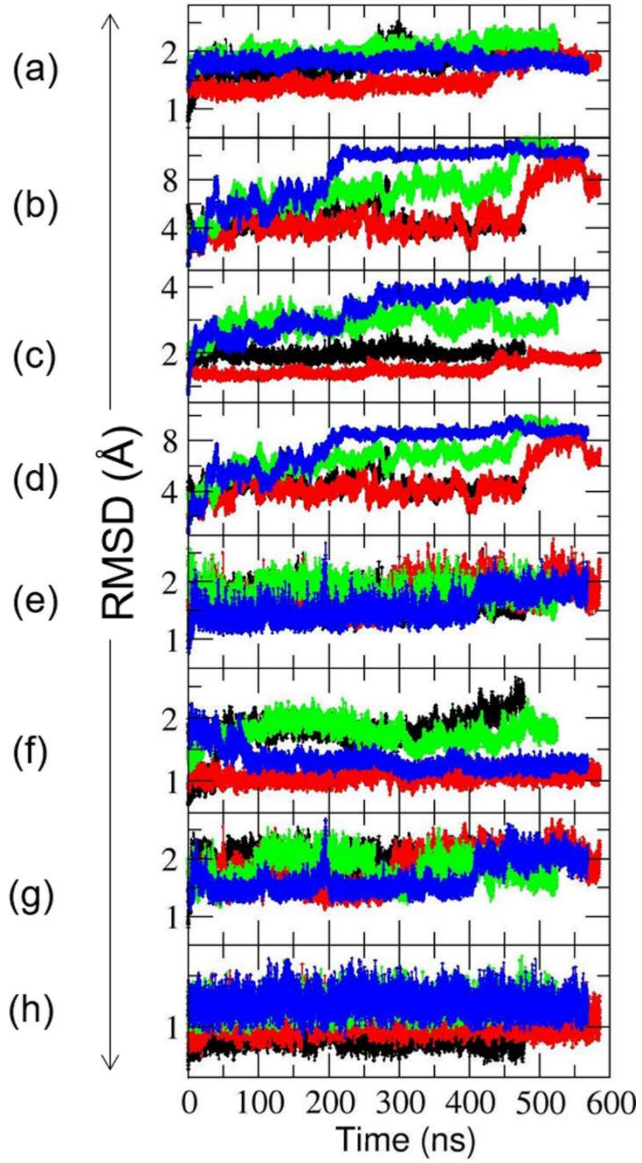


**Figure S4:** Observed $C_{\alpha}$-RMSD changes during the simulations of membrane-bound CYP 1A1-CPR complexes. Cα atom root mean squared deviation (Cα-RMSD) of (a) the globular domain of CYP 1A1, (b) the globular domain of CPR, (c) the globular domain of CYP and FMN domain of CPR, (d) the FMN and FAD domains of CPR. (e) the FAD and NADP domains of CPR, (f) the FMN domain only, (g) the FAD domain only and (h) the NADP domain only. The RMSDs were calculated with respect to the initial energy minimized encounter complexes generated by BD simulation. **Color scheme:** C2: Black; C2: Red; D2: Green; D2': Blue.

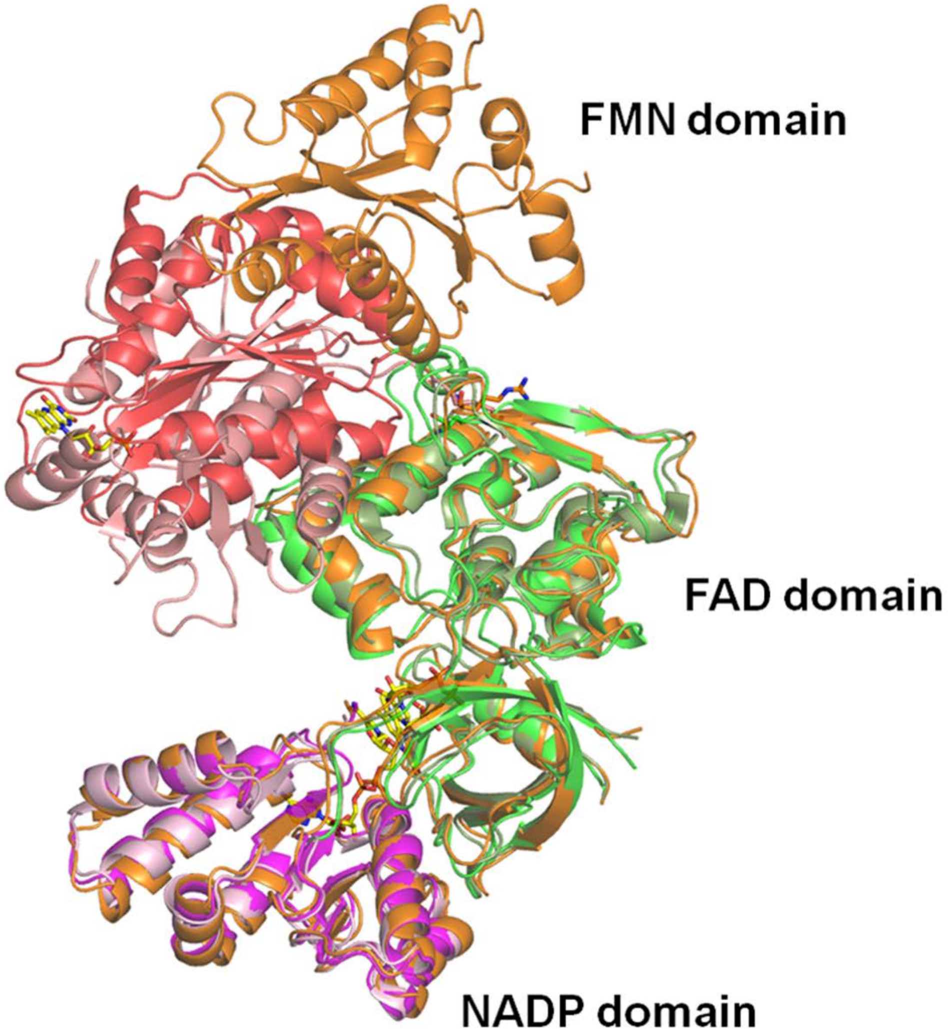


**Figure S5:** Closure of the CPR structure during simulations of the CYP 1A1-CPR complex in a membrane compared to crystallographic structures of CPR. The structures of chain A (semi-closed; NADP domain: pale pink, FAD domain: pale green, FMN domain: salmon) and chain B (open; brown, energy minimized) in the crystal structure (PDB ID: 3ES9) as well as the structure obtained after ~500 ns of the D2' simulation starting from the chain B conformation (NADP domain: magenta, FAD domain: green, FMN domain: red) are shown in cartoon representation after superposition of the Cα atoms of the NADP and FAD domains.

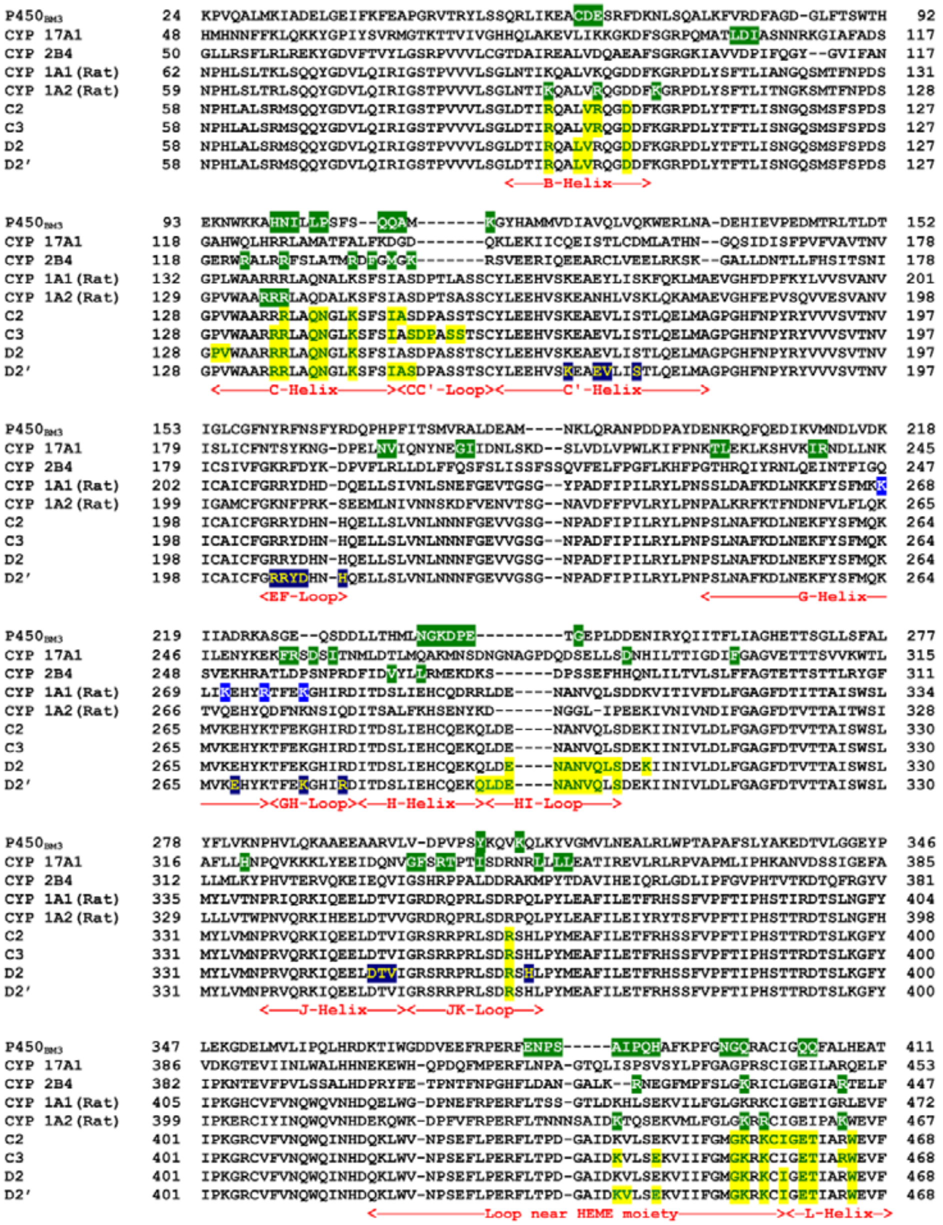


**Figure S6:** Multiple sequence alignment of *Bacillus megaterium* CYP BM3 ($P450_{BM3}$), human CYP 17A1, rabbit CYP 2B4, rat CYP 1A1, rat CYP 1A2, and human CYP 1A1 (lower four rows). Green highlighted white residues are residues for which there is site-directed mutagenesis or structural evidence (CYP1A2(rat)-CPR,[32] CYP17A1-CPR,[33,34] CYP2B4-CPR,[35] CYP1A1(rat)-CPR,[36–38] $P450_{BM3}$[39]) indicating that they are at the interface with the CPR FMN domain. Note that in CYP17A1, all these residues are on the distal side except G337, F338, R340, T341, I344, L350, L352 and L353, and are indicated by NMR to be affected by FMN domain binding.[34] Blue highlighted white residues are distal site residues affected by CPR binding according to site-directed mutagenesis data. Yellow highlighted green residues are residues at the interface with the CPR FMN domains and navy highlighted yellow residues are residues at the interface with the CPR NADP domains in the four model complexes. The secondary structure of the CPR binding interface of human CYP 1A1 is given under each row.

Multiple sequence alignment was performed using the PROMALS3D webserver (http://prodata.swmed.edu/).[40]

```
Aromatase        377  ALEDDVIDGYPVKKGTNIILNIGRMHRL-EFFPKPNEFTLENFAK------NVPYRYFQPFGFGPRGCAG
CYP 3A4          377  CKKDVEINGMFIPKGVVVMIPSYALHRDPKYWTEPEKFLPERFSKKNKD--NIDPYIYTPFGSGPRNCIG
CYP 1A1          390  TTRDTSLKGFYIPKGRCVFVNQWQINHDQKLWVNPSEFLPERFLTPDGAIDKVLSEKVIIFGMGKRKCIG

Aromatase        440  KYIAMVMMKAILVTLLRRFHVKTLQGQCVESIQKIHDLSLHPDETKNMLEMIFTPRNSDRCLEH  503
CYP 3A4          445  MRFALMNMKLALIRVLQNFSFKPCKETQIP-LKLSLGGLLQPE---KPVVLKVESRDGTVSGA-  503
CYP 1A1          460  ETIARWEVFLFLAILLQRVEFSVPLGVKVD-MTPIYGLTMKHA---CCEHFQMQLRS-------  512
```

**Figure S7:** Part of the multiple sequence alignment of human aromatase, CYP 3A4 and CYP 1A1. Yellow highlighting shows predicted electron transfer pathways between redox centers. Multiple sequence alignment was performed using the PROMALS3D webserver (http://prodata.swmed.edu/).[40]

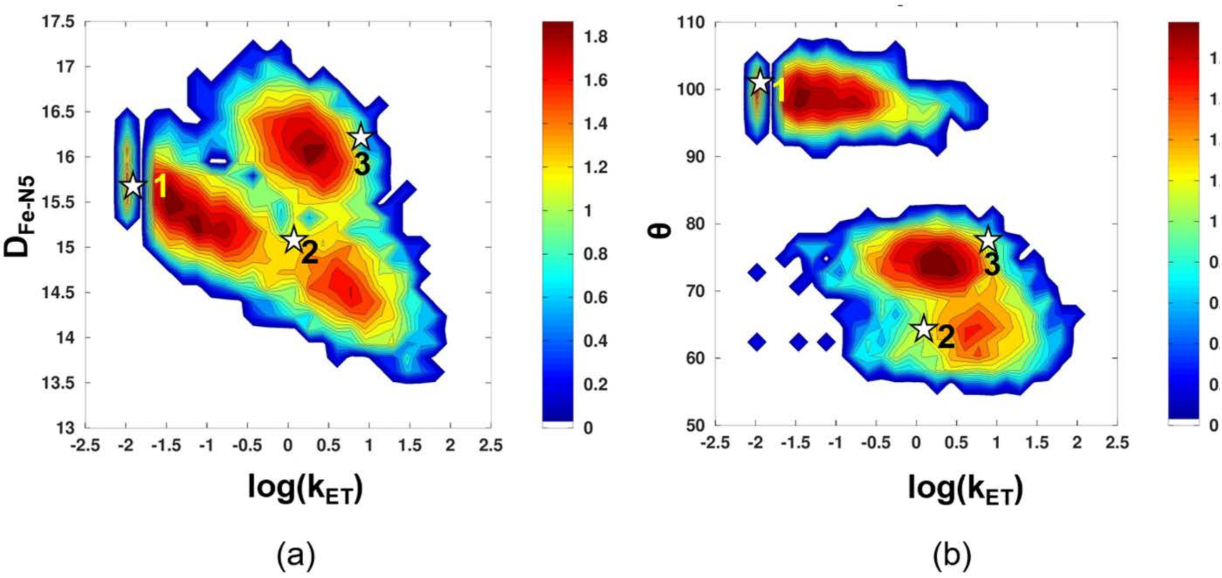


**Figure S8:** 2D histogram plots of (a) the distance between the redox centers ($D_{Fe\text{-}N5}$) and (b) the angle θ between the α1-helix of the FMN domain of CPR and the C-helix of CYP 1A1 vs $\log(k_{ET})$. Stars indicate the representative complexes from the last snapshots of the simulations: (1) C3, (2) D2 and (3) D2’. The atomic coordinates of these complexes are provided in the Supplementary Information.

### Supplementary Tables

**Table S1:** Properties of the eight most energetically favorable unique clusters of diffusional encounter complexes of CPR and CYP 1A1 obtained by BD rigid-body docking with SDA. The selected clusters were required to have more than 100 structures of complexes, (CLsize > 100). For details of the selection criteria, see Methods and Supplementary Methods.

| **Complexes** | **ClSize*** | **ClFSize#** | **ReprE$** | **ClAE&** |
|---|---|---|---|---|
| C3 | 368 | 37490 | -33.8 | -35.5 |
| D3 | 239 | 13572 | -33.0 | -33.3 |
| D2 | 261 | 13084 | -28.1 | -27.4 |
| A6 | 237 | 15495 | -27.9 | -26.2 |
| C2 | 485 | 26602 | -27.7 | -27.5 |
| B7 | 198 | 14843 | -27.6 | -27.6 |
| A4 | 248 | 14737 | -27.5 | -28.7 |
| B2 | 424 | 28458 | -27.3 | -25.6 |

*ClSize: Number of saved structures of complexes in the cluster
#ClFSize: Total number of complexes in the cluster
$ReprE: Total interaction energy in kT of the representative structure for the cluster
&ClAE: Average total energy in kT of all saved structures in the cluster weighted by the number of occurrences of each structure.

**Table S2:** Variations in the interface residues of CYP 1A1 and CPR observed in the six selected BD rigid-body docked encounter complexes (A4, B7, C2, C3, D2 and D3) and identified after contact analysis of the soluble simulations of these encounter complexes and the membrane simulations based on three encounter complexes (C2, C3, D2 and D2'). The occupancy of the listed CYP-FMN domain interactions was 70% or higher in the MD simulations.

| Complex | CYP interface | CPR interface |
|---|---|---|
| A4 | B-Helix: **R93**, **V97**, **R98**<br>C-Helix: R136, **K143**, *I147*<br>CC'-Loop: *A148*, *S149*, *S154*<br>JK-Loop: *R362*, *S363*<br>Loop near HEME: **K441**, *V442*, **E445**, *M452*, *G453*, *K454*, **K456**<br>L-Helix: *T461*, *W465* | Lβ1-Loop: *Q87*, *T88*, *T90*<br>Lβ2-Loop: *E115*, *E116*<br>Lβ3-Loop: *G141*, **E142**, *D147*<br>α3-Helix: **Q150**, D154<br>Lβ4-Loop: *T177*, **Y178**, **E179**, H180<br>Lβ5-Loop: *D208* |
| B7 | B-Helix: R93, **V97**, R98, D101<br>C-Helix: *R135*, *R136*, *Q139*, *L142*, *K143*, *I147*<br>JK-Loop: *R362*, *S363*, *H364*<br>Loop near HEME: **K441**, **V442**, *E445*, *M452*, **G453**, **K454**, *K456*, *I458*<br>L-Helix: *E460*, **T461**, *R464*, **W465** | Lβ1-Loop: *Q87*<br>Lβ3-Loop: **Y140**, *G141*, **E142**, **D144**, *P145*, *T146*, *D147*<br>α3-Helix: *Q157*<br>Lβ4-Loop: N175, *K176*, **T177**, **Y178**, *E179*, *H180*<br>Lβ5-Loop: D208, **D209** |
| C2 | B-Helix: **R93**, **V97**, **R98,** D101<br>C-Helix: R136, Q139, **N140**, **K143**, **I147**<br>CC'-Loop: A148<br>JK-Loop: R362<br>Loop near HEME: *V442*, *E445*, G453, **K454**, **K456**, C457, I458<br>L-Helix: G459, *E460*, **T461**, **W465** | Lβ1-Loop: Q87, T88<br>Lβ2-Loop: *E115*, L119<br>Lβ3-Loop: Y140, G141, **E142**, **D144**, **D147**<br>α3-Helix: **Q150**, **D151**, **Y153**, **D154**, W155, **Q157**, E158<br>Lβ4-Loop: N175, **T177**, **Y178**, **E179**, **H180** |
| C3 | B-Helix: R93, L96, **V97**, **R98**, **D101**<br>C-Helix: *W131*, *A132*, **R135**, **R136**, **Q139**, **N140**, **K143**, I447<br>CC'-Loop: S149, D150, P151, S153, S154<br>JK-Loop: R362<br>Loop near HEME: **K441**, **E445**, *M452*, **G453**, **K454**, *K456*<br>L-Helix: **E460**, **T461**, R464, **W465** | Lβ1-Loop: **Q87**, **T88**, G89 , T90<br>α1-Helix: E92<br>Lβ2-Loop: **E115**<br>Lβ3-Loop: Y140, G141, **E142**, **D144**, **D147**<br>α3-Helix: **Q150**, **D151**, **D154**, *Q157*, E158<br>Lβ4-Loop: **T177**, **Y178**, *E179*, **H180**<br>Lβ5-Loop: D208, D209, G210, N211 |

| | | |
|---|---|---|
| D2 | B-Helix: *R93*, L96, *V97*, D101<br>C-Helix: P129, V130, *R135*, **R136**, Q139, N140, **K143**, S144, *I147*<br>HI-Loop: E296, N297, A298, N299, V300, Q301, L302, S303<br>I-Helix: K306<br>J-Helix: D348, T349, V350<br>JK-Loop: S360, **R362**, **S363**, H364<br>Loop near HEME: **K441**, **E445**, *I448*, *M452*, **G453**, **K454**, *K456*, I458<br>L-Helix: **E460**, **T461**, **R464**, **W465** | β1-Sheet: Y84<br>Lβ1-Loop: S86, **Q87**, **T88**, G89, T90<br>α1-Helix: E92<br>Lβ2-Loop: E115<br>Lβ3-Loop: **Y140**, **G141**, **E142**, G143, **D144**, D147, N148<br>α3-Helix: D151<br>Lβ4-Loop: **N175**, **K176**, **T177**, **Y178**, **E179**, *H180*<br>Lβ5-Loop: **D208**, **D209**, **G210**, N211<br>NADP domain: *K664*, *T667*, *K668* |
| D2' | B-Helix: *R93*, L96, *V97*, R98, D101<br>C-Helix: *R135*, **R136**, Q139, N140, **K143**, S144, *I147*<br>CC'-Loop: A148, S149<br>C'-Helix: K165, E168, V169, S172<br>EF-Loop: R205, R206, Y207, D208 H211<br>G-Helix: E268<br>GH-Loop: K275 R279<br>HI-Loop: Q293, L294, D295, E296, N297, A298, N299, V300, Q301, S303<br>JK-Loop: S360, **R362**, **S363**<br>Loop near HEME: **K441**, V442, **E445**, *I448*, *M452*, **G453**, **K454**, *K456*, I458<br>L-Helix: **E460**, **T461**, **R464**, **W465** | β1-Sheet: R78<br>Lβ1-Loop: **Q87**, **T88**, G89, T90<br>α1-Helix: E92, E93, N96<br>Lβ2-Loop: S111, E115<br>Lβ3-Loop: **Y140**, **G141**, **E142**, **D144**, D147<br>α3-Helix: D151<br>Lβ4-Loop: **N175**, **K176**, **T177**, **Y178**, **E179**, *H180*<br>Lβ5-Loop: **D208**, **D209**, **G210**, N211<br>NADP domain: E654, *H655*, A656, Q657, V659, D660, *K663*, K664, *M666*, T667 |
| D3 | B-Helix: ***R93***, ***V97***, *R98*, **D101**<br>C-Helix: **R136**, **Q139**, *N140*, **K143**, *I147*<br>JK-Loop: *S363*<br>Loop near HEME: **K441**, **V442**, **E445**, K446, M452, G453, **K454**, *K456*<br>L-Helix: *E460*, *T461* | Lβ1-Loop: *S86*, **Q87**<br>Lβ2-Loop: **E115**<br>Lβ3-Loop: *G141*, **E142**, **D144**, **D147**<br>α3-Helix: Q150, **D151**, Y153, **D154**, *Q157*<br>Lβ4-Loop: **T177**, **Y178**, **E179**, **H180** |

**Color Scheme:** Blue font: Residues present at the interface at all simulation stages (bold); in soluble and membrane simulations (standard font), and in BD docking and membrane simulations (italic) and only in membrane simulations (underlined). Black font: Residues present at the interface after BD docking and soluble simulations (bold), only after BD docking (italic), only in soluble simulations (standard font). The NADP domain interface residues which are 70% or more than contact occupancy are highlighted in blue background and standard font, whereas, the NADP domain residues which have contact occupancy less than 70% are in blue background and italic font.

**Table S3:** Structural rearrangement during the soluble MD simulations of six complexes of the CYP 1A1 globular domain and the CPR FMN domain in aqueous solution. The $C_\alpha$-RMSD values of the globular domain of CYP 1A1 and the FMN domain of CPR show that there is little variation in the internal structure of the individual domains. The movement of the FMN domain with respect to CYP is shown in the last column by the $C_\alpha$-RMSD of the FMN domain when the complex was superimposed on the first frame using the CYP $C_\alpha$ atoms only for each snapshot analyzed. All values were computed for the last 20 ns of the trajectories.

| Complex | Simulation length (ns) | RMSD $_{CYP\ domain}$ (Å) | RMSD $_{FMN\ domain}$ (Å) | Movement of FMN domain (Å) |
|---|---|---|---|---|
| A4 | 98.1 | 2.0±0.1 | 1.4±0.1 | 12.4±0.8 |
| B7 | 135.5 | 2.0±0.1 | 1.3±0.1 | 17.1±3.4 |
| C2 | 122.6 | 2.0±0.1 | 1.2±0.1 | 9.3±1.0 |
| C3 | 100.4 | 2.6±0.1 | 1.2±0.1 | 3.1±0.5 |
| D2 | 48.1 | 2.1±0.1 | 1.4±0.1 | 4.2±0.6 |
| D3 | 92.7 | 2.0±0.2 | 1.3±0.1 | 4.7±0.9 |

**Table S4:** Parameters characterizing the positioning and configuration of the CYP-CPR complexes with respect to the membrane during the simulations based on encounter complexes C2, C3 and D2. D2' is a replica simulation based on encounter complex D2.

| **Parameters *** | **Initial value (for start frame)**[+] | | | **CYP - membrane simulation in absence of CPR (last frame)** | **Final value (average over last 50 ns for snapshots at 100 ps intervals)** | | | |
|---|---|---|---|---|---|---|---|---|
| | **C2** | **C3** | **D2** | | **C2** | **C3** | **D2** | **D2'** |
| Angle α (°) | 95.2 | 93.8 | 94.0 | 96.0 | 70.3±2.8 | 83.4±3.6 | 67.5±2.4 | 72.4±2.0 |
| Angle β (°) | 139.9 | 140.1 | 138.7 | 135.3 | 109.7 ±3.2 | 127.1 ±4.3 | 99.4±2.6 | 105.5 ±2.0 |
| Angle Heme tilt (°) | 71.4 | 75.5 | 71.1 | 67.6 | 41.2±4.4 | 59.1±5.3 | 36.1±3.8 | 38.6±3.1 |
| $\theta$ (°) | 94.0 | 96.4 | 62.1 | --- | 95.7±4.0 | 99.1±1.8 | 62.0±3.1 | 75.1±2.6 |
| $TM_{CYP}$ helix tilt ($\gamma_{CYP}$ °) | 27.9 | 27.3 | 26.1 | 28.3 | 14.2±5.4 | 46.8±3.3 | 25.6±3.9 | 45.6±3.6 |
| $TM_{CPR}$ helix tilt ($\gamma_{CPR}$ °) | 21.1 | 15.3 | 16.0 | --- | 30.9±3.5 | 37.9±4.7 | 18.9±4.1 | 13.7±4.9 |
| $D_{CYP\text{-}mem}$ (Å) | 42.1 | 41.8 | 42.1 | 42.1 | 49.9±1.7 | 44.8±2.7 | 49.6±2.1 | 49.7±2.0 |
| $D_{FMN\ domain\text{-}mem}$ (Å) | 70.5 | 70.2 | 76.0 | --- | 61.9±1.3 | 66.6±1.3 | 62.3±1.3 | 61.1±2.1 |
| $D_{FAD\ domain\text{-}mem}$ (Å) | 99.2 | 83.5 | 97.9 | --- | 46.4±1.8 | 60.2±2.6 | 97.2±1.2 | 81.3±2.1 |
| $D_{NADP\ domain\text{-}mem}$ (Å) | 112.3 | 87.5 | 88.4 | --- | 47.2±2.6 | 67.4±4.2 | 99.4±2.2 | 75.4±2.2 |
| $D_{FG\text{-}mem}$ (Å) | 21.0 | 22.0 | 20.6 | 20.8 | 23.9±1.7 | 21.4±2.2 | 25.6±2.3 | 24.1±2.0 |
| $D_{linker\_mem}$ (Å) | 60.7 | 58.5 | 76.4 | --- | 46.4±1.6 | 53.9±3.7 | 51.8±1.5 | 43.6±2.9 |
| $D_{Fe\text{-}N5}$ (Å) | 17.0 | 15.3 | 14.5 | --- | 15.6±0.3 | 15.5±0.4 | 14.7±0.4 | 16.3±0.3 |
| $D_{CYP\text{-}FMN\ domain}$ (Å) | 36.2 | 34.2 | 36.2 | --- | 34.6±0.2 | 33.7±0.2 | 35.6±0.4 | 33.0±0.2 |
| $D_{CYP\text{-}FAD\ domain}$ (Å) | 79.3 | 69.9 | 68.5 | --- | 65.6±1.2 | 60.0±1.9 | 57.0±0.7 | 54.1±0.4 |
| $D_{CYP\text{-}NADP\ domain}$ (Å) | 95.9 | 84.8 | 78.4 | --- | 80.9±1.6 | 88.6±3.9 | 50.9±0.7 | 45.5±0.3 |
| $D_{FAD\text{-}NADP\ domain}$ (Å) [%] | 34.3 | 34.3 | 34.1 | --- | 34.2±0.3 | 34.8±0.3 | 33.9±0.3 | 33.9±0.3 |
| $D_{FMN\text{-}NADP\ domain}$ (Å) [%] | 71.5 | 71.5 | 71.6 | --- | 68.5±1.1 | 75.1±0.9 | 49.5±0.8 | 53.9±0.5 |
| $D_{FMN\text{-}FAD\ domain}$ (Å) [%] | 47.7 | 47.5 | 47.8 | --- | 43.4±0.4 | 42.8±0.5 | 37.0±0.8 | 39.6±0.5 |
| $D_{TM\text{-}TM}$ (Å) | 32.0 | 33.6 | 32.8 | --- | 39.6±1.4 | 22.2±1.2 | 29.1±0.8 | 38.8±1.5 |
| ΔASA[#] (Å$^2$) | 1790 | 2377 | 1773 | --- | 2000 ±135 | 2453 ±158 | 1817 ±229 | 2480 ±142 |

*Angles α, β, γ and the heme tilt angle of CYP 1A1 as well as angle $\theta$, between the α1 helix of the FMN domain and the C helix of CYP 1A1, are defined in Fig. 2. α, β and γ are the angles between the z-axis and the vectors v1, v2 and v3, respectively, defined as follows. v1: along the I-helix, connecting the centers of the first (residues 307-310) and last (331-336) helical turns; v2: orthogonal to v1, connecting one helical turn in the C-helix (129-133) and one in the F-helix (224-228); v3: along the TM-helix connecting the centers of the first (11-14) and last (21-24) helical turns. The heme tilt angle is the angle between the heme plane and the z-axis perpendicular to the membrane plane. The angle θ was calculated between vectors along the C-helix of CYP 1A1, connecting the centers of the first three (129-132) and last three (143-146) residues, and the α1-helix of the FMN domain, connecting the centers of the first three (90-93) and the last three (102-105) residues. The $TM_{CPR}$ tilt angle was calculated between the CPR TM-helix, connecting the centers of the first four (25-29), last four (36-40) residues, and the z-axis. Angles were calculated by computing the dot product and length of the two respective vectors and the angle was obtained using the cosine formula.

Distances (Å): center-to-center distance between CYP (residues 51-511) and membrane ($D_{CYP\text{-}mem}$), FMN domain (residues 66-230) and membrane ($D_{FMN\ domain\text{-}mem}$), FAD domain (residues 247-514) and membrane ($D_{FAD\ domain\text{-}mem}$), NADP domain (residues 518-677) and membrane ($D_{NADP\ domain\text{-}mem}$), FG region of CYP (residues 229-245) and membrane ($D_{FG\text{-}mem}$), linker of CPR (residue N of E66) and membrane ($D_{linker\text{-}mem}$), redox center distance ($D_{Fe\text{-}N5}$), center-to-center distances between CYP and FMN domain ($D_{CYP\text{-}FMN\ domain}$), CYP and FAD domain ($D_{CYP\text{-}FAD\ domain}$), CYP and NADP domain ($D_{CYP\text{-}NADP\ domain}$), FAD and NADP domain ($D_{FAD\text{-}NADP\ domain}$), and $TM_{CYP}$ (residues 7-26) and $TM_{CPR}$ (residues 25-44) ($D_{TM\text{-}TM}$). The values for the membrane-bound CYP 1A1 in the absence of CPR are given for comparison.

[+]The starting orientations of the four complexes differ with respect to the membrane-bound CYP 1A1 system (in the absence of CPR) due to the different degree of superimposition of the globular domain of CYP 1A1 on the membrane-bound CYP 1A1 simulated in the absence of CPR and the CYP 1A1-FMN domain of CPR complexes obtained from soluble MD simulations. In the soluble simulations, there was a deviation of 2-2.6 Å Cα-RMSD of the globular domain of CYP 1A1 with respect to the initial structure.

[%] In the crystal structure of CPR, the interdomain distances have the following values in Å in chain A/B, respectively: $D_{FAD\text{-}NADP\ domain}$ = 34.3/33.8; $D_{FMN\text{-}NADP\ domain}$ = 45.9/70.4; $D_{FMN\text{-}FAD\ domain}$ =35.5/46.7. The simulations were started from a conformation close to that of chain B. During the simulations, the CPR tended to close and approach the conformation of chain A.

[#]ΔASA is the interface contact area of globular domain of CYP and FMN domain of CPR.

(26) Marcus, R. A. Electrostatic Free Energy and Other Properties of States Having

Nonequilibrium Polarization. I. *J. Chem. Phys.* **1956**, *24* (5), 979–989.
